# Uncovering active modulators of native macroautophagy through novel high-content screens

**DOI:** 10.1101/756973

**Authors:** N Safren, EM Tank, N Santoro, SJ Barmada

**Affiliations:** University of Michigan, Department of Neurology, Ann Arbor MI; University of Michigan, Center for Chemical Genomics, Life Sciences Institute

## Abstract

Autophagy is an evolutionarily conserved pathway mediating the breakdown of cellular proteins and organelles. Emphasizing its pivotal nature, autophagy dysfunction contributes to many diseases; nevertheless, development of effective autophagy modulating drugs is hampered by fundamental deficiencies in available methods for measuring autophagic activity, or flux. To overcome these limitations, we introduced the photoconvertible protein Dendra2 into the *MAP1LC3B* locus of human cells via CRISPR/Cas9 genome editing, enabling accurate and sensitive assessments of autophagy in living cells by optical pulse labeling. High-content screening of 1,500 tool compounds provided construct validity for the assay and uncovered many new autophagy modulators. In an expanded screen of 24,000 diverse compounds, we identified additional hits with profound effects on autophagy. Further, the autophagy activator NVP-BEZ235 exhibited significant neuroprotective properties in a neurodegenerative disease model. These studies confirm the utility of the Dendra2-LC3 assay, while simultaneously highlighting new autophagy-modulating compounds that display promising therapeutic effects.

## Introduction

Macroautophagy (hereafter referred to as autophagy) is an essential pathway for protein homeostasis whereby cytoplasmic proteins and organelles are delivered to lysosomes for degradation^1^. Through the coordinated action of a series of autophagy-related (ATG) proteins and cargo receptors including p62/SQSTM1, NBR1, and optinuerin^2^, substrates are sequestered within double membrane vesicles called autophagosomes. Autophagosomes mature as they traffic along microtubules and eventually fuse with lysosomes to form autolysosomes, wherein hydrolases degrade autophagic cargo. The protein LC3 (ATG8) is an obligate component of autophagosome membranes and is itself degraded within autolysosomes. For these reasons, it often serves as both a marker of autophagosomes and as a representative autophagy substrate^3^.

Underscoring the critical requirement of autophagy in cellular homeostasis, deletion of core autophagy genes in mice results in embryonic lethality^4, 5, 6^. Accordingly, dysfunctional autophagy is linked to a wide spectrum of human disease including neurodegeneration, cancer, metabolic disorders, infectious and cardiovascular diseases^7^. Often these conditions involve deficiencies in one or more steps of autophagy, resulting in impaired clearance of potentially toxic cellular components, and/or a failure to replenish amino acids required for anabolic processes. In these instances, enhancing the rate of autophagic cargo clearance, commonly referred to as flux, would be beneficial. Conversely, autophagy can promote tumor progression and resistance to chemotherapy for some cancers^8, 9, 10, 11^. Here, autophagy inhibition may represent a more apt therapeutic strategy^7^.

Autophagy is of particular importance in the central nervous system (CNS). Deletion of essential autophagy genes within the CNS of mice leads to progressive neurodegeneration marked by accumulation of protein aggregates^12, 13, 14, 15, 16^. Defective autophagy is a common feature of many neurodegenerative diseases including Alzheimer’s disease^17, 18^, Parkinson’s disease^19, 20, 21, 22, 23^, polyglutamine disorders^24, 25, 26^, amyotrophic lateral sclerosis (ALS)^27^ and frontotemporal dementia (FTD)^28, 29, 30, 31^. Moreover, mutations in several autophagy related proteins including p62/SQSTM1^32^, optineurin^33^, C9ORF72^34, 35^, TBK1^36^, and UBQLN2^37^ results in familial ALS and FTD.

Due to its broad therapeutic potential, autophagy modulation has received considerable attention as a target for drug development^7^. Nevertheless, these efforts have thus far failed to translate into effective therapies for patients. This is in part due to the intrinsic difficulties in measuring autophagic flux, and consequent inability of many conventional and widely used autophagy assays to accurately estimate flux^3^. One prominent limitation of these assays is an implicit reliance on the steady-state abundance of pathway intermediates such as LC3-II, the lipidated isoform of LC3. Due to the dynamic nature of autophagy, changes in such intermediates may equally reflect increased autophagy induction or late-stage inhibition of autophagsome clearance; although discriminating among these mechanisms is crucial for drug development, many assays are effectively unable to do so. While lysosomal inhibitors such as bafilomycin-A1 can be used to isolate autophagy induction from inhibition, this approach obscures estimates of substrate clearance, perhaps the most relevant measure of autophagic flux. Bafilomycin-A1 and related compounds are also inherently toxic, further confounding flux measurements^38, 39^. Yet another common shortcoming is an inherent reliance on static “snapshots” of separate cellular populations that cannot be followed prospectively or longitudinally due to the need for cell lysis and measurement of pathway intermediates.

We previously developed a technique called optical pulse labeling (OPL), enabling non-invasive measurements of autophagic flux in living cells without the need for lysosomal inhibition^40^. In this technique, LC3 is labeled with the photoconvertible protein Dendra2^41^. Upon exposure to blue (405nm) light, Dendra2 fluorescence irreversibly shifts from green to red. Since the generation of red-fluorescent Dendra2-LC3 is limited by blue light, LC3 turnover can be determined independent of new protein synthesis by tracking the time-dependent reduction in red fluorescence following a brief pulse of blue light (Fig. 1A). LC3 is an autophagy substrate, and therefore its degradation kinetics serves as a faithful proxy for estimates of autophagic flux. While OPL offered several advantages over conventional assays, it was nonetheless limited by its reliance on protein overexpression; in effect, Dendra2-LC3 overexpression floods the pathway under investigation with an obligate substrate. Burdening the cell with non-physiological concentrations of substrate might artificially prolong flux estimates, or conversely enhance flux via regulatory feedback mechanisms. Moreover, because autophagy regulation is intricately tied to amino acid availability^42, 43^ and the ubiquitin proteasome system^44^, any perturbations to these pathways brought on by protein overexpression may further confound measurements of flux.

**Figure 1:**
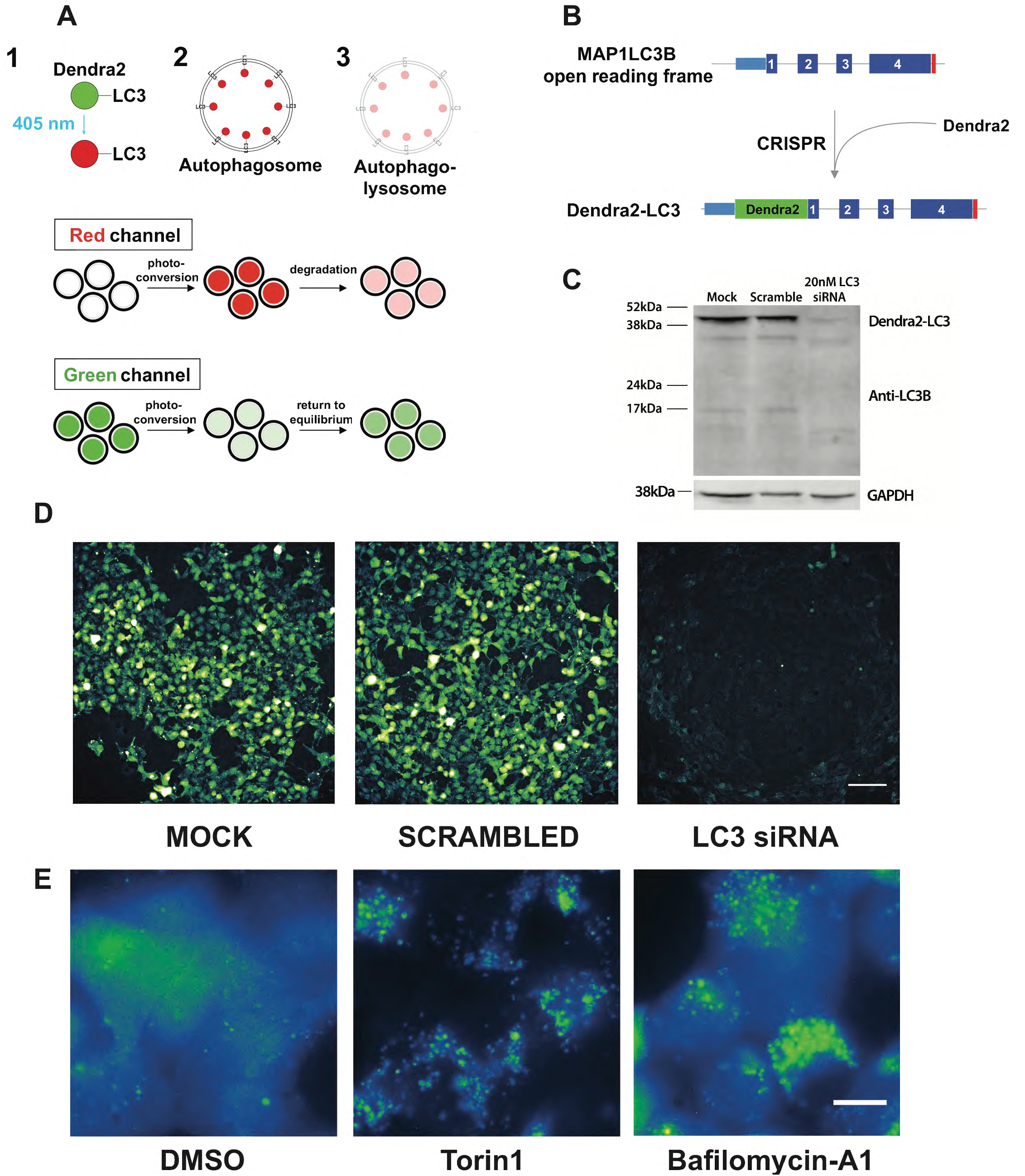
Creation of a stable cell line serving as a reporter for autophagic flux. (**A**) Illustration depicting the use of optical pulse labeling (OPL) to measure autophagic flux. Dendra2 is a photoconvertible protein that upon exposure to 405nm light irreversibly shifts its fluorescence from green to red. Photoconverted Dendra2-LC3 is degraded in lysosomes, resulting in a drop in red fluorescence intensity. The time dependent decay of red signal serves as an estimate of autophagic flux independent of new (green) LC3-Dendra2 synthesis. (**B**) Schematic for tagging native LC3 using CRISPR/Cas9 genome editing. In HEK293T cells, the Dendra2 ORF was introduced into the *MAP1LC3B* locus upstream of exon 1 creating an N-terminal fusion protein upon translation. (**C**) Western blot confirming the successful labeling of LC3 with Dendra2. Dendra2-LC3 HEK293T cells were treated with 20nM siRNA targeting LC3 or scrambled siRNA. Lysates were collected after 48h and immunoblotted with an LC3 antibody, demonstrating the Dendra2-LC3 fusion protein running at the expected MW of 43kDa that disappears upon siRNA-mediated knockdown of LC3. GAPDH serves as a loading control. (**D**) Dendra2-LC3 reporter line imaged in GFP channel 48h after application of siRNA. Scale bar = 100µm (**E**) Dendra2-LC3 cells imaged 6 hours after treatment with vehicle, 1µM Torin1, and 20nM Bafilomycin-A1. Scale bar =10µm

To overcome these drawbacks, we labeled *native* LC3 with Dendra2 using CRISPR/Cas9 editing, producing a novel autophagy reporter cell line capable of assaying flux in living cells without the need for drug treatment, protein overexpression or measurements of pathway intermediates, thus establishing a faithful reporter of endogenous autophagy activity unadulterated by exogenous manipulations. Leveraging this cell line for its unique perspective on autophagy and the opportunities it presents, we adapted Dendra2-LC3 cells for conducting high-content screens and identified several new and active autophagy modulators with promising therapeutic properties.

## Results

### Creation of a novel reporter of autophagic flux

We employed CRISPR/Cas9 genome editing to label native LC3 by introducing Dendra2 into the *MAP1LC3B* locus (encoding LC3) of human embryonic kidney (HEK) cells^45^ (Fig. 1B). To minimize the risk of undesired insertions/deletions via non-homologous end-joining we used a dual-nickase strategy^46^, in which Cas9(D10A) was expressed along with two single-guide RNAs (sgRNAs) targeting sequences immediately upstream and downstream of the *MAP1LC3B* start codon. Unlike wild-type Cas9, Cas9(D10A) induces single-stranded nicks rather than double-stranded breaks in the DNA, limiting recombination to the region marked by the sgRNAs. In addition, a vector containing the Dendra2 open reading frame (ORF) flanked by 400bp of homologous sequence 5’ and 3’ to the *MAP1LC3B* start codon was introduced to facilitate homology directed repair (HDR), thereby creating a sequence encoding Dendra2 fused to the N-terminus of LC3. Positive cells were selected based on Dendra2 fluorescence and enriched by sequential passaging until a homogenous population was achieved. Western blotting confirmed the successful insertion of the Dendra2 ORF into the desired locus (Fig. 1C). Transfection with siRNA targeting LC3 substantially reduced both Dendra-LC3 protein levels and native GFP fluorescence, providing further validation of successful on-target CRISPR editing (Fig. 1C,D).

In untreated cells, Dendra2-LC3 fluorescence was diffusely distributed, matching the predicted localization of the non-lipidated, cytosolic LC3-I isoform (Fig. 1E, Supplemental movie 1). Treatment with the potent autophagy inducer Torin1^47^ elicited the formation of visible fluorescent puncta and reduced the intensity of diffuse Dendra2-LC3 (Fig. 1E, Supplemental movie 2), reflecting the incorporation of Dendra2-LC3 into autophagosome membranes. In agreement with previous studies of autophagosome dynamics^26, 48^, live cell imaging revealed that a subset of Dendra2-LC3 puncta were highly mobile (Supplemental movie 2). As expected, inhibiting the clearance of autophagosomes via treatment with the lysosomal V-ATPase inhibitor bafilomycin-A1 lead to the accumulation of large bright puncta without an accompanying decrease in diffuse Dendra2-LC3 fluorescence (Supplemental movie 3). Together these data confirm that Dendra2-tagged version of LC3 behaves as expected in modified HEK293T cells^40^, and that these cells can be used to visualize autophagy modulation by a variety of stimuli.

### Development and validation of an autophagic flux assay

In these cells, endogenous Dendra2-LC3 could be efficiently photoconverted with minimal toxicity using 4s pulses of 405nm light, producing a strong red signal (Fig. 2A) concurrent with a substantial reduction in green fluorescence. Using time-lapsed microscopy, we measured fluorescence intensity in both the red (TRITC) and green (GFP) channels at regular intervals over the span of 13.5h. In vehicle-treated cells red fluorescence decayed with a half-life of approximately 7.5h. Treatment with Torin1 significantly accelerated this decay, reducing Dendra2-LC3 half-life ∼3-fold to 2.5h. In contrast, bafilomycin-A1 completely stabilized Dendra2-LC3 and blocked the Torin1-induced reduction in Dendra2-LC3 half-life (Fig. 2A, B). Thus, endogenous Dendra2-LC3 flux measured by OPL responds appropriately to bidirectional modulation of autophagy.

**Figure 2:**
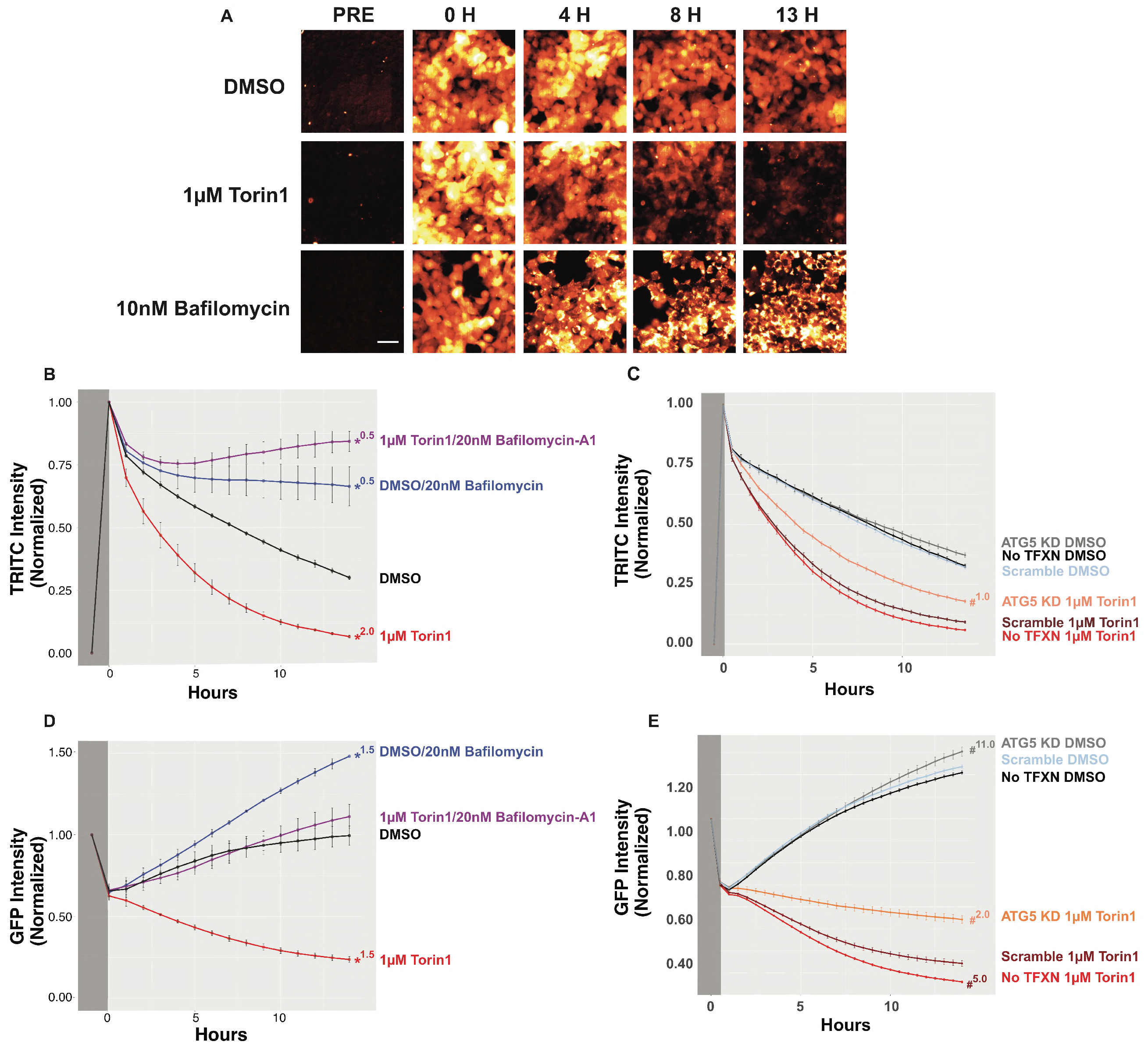
Time dependent decay of Dendra2-LC3 serves as an accurate measure of autophagic flux. (**A**) Dendra2-LC3 HEK293T cells were imaged prior to photoconversion to measure background RFP intensity. Immediately following photoconversion, cells were treated with DMSO, 1µM Torin1, or 10nM Bafilomycin-A1 and imaged at the indicated times. Images are pseudocolored to better highlight intensity differences. Scale bar = 50µm (**B-E**) Time dependent changes in photoconverted Dendra2-LC3 fluorescence in the RFP **(B, C)** and GFP (**D, E**) channels. Intensity measurements were obtained prior to (dark grey) and following photoconversion (light grey) and normalized. For RFP measurements, the background intensity prior to conversion was set to 0 and the post-conversion value to 1. GFP values are scaled to the pre-conversion intensity. Error bars represent SEM from 3 replicate experiments. (**B**) Treatment with 1µM Torin1 accelerates Dendra2-LC3 decay, reflecting enhanced autophagic degradation of the reporter, while treatment with bafilomycin-A1 stabilizes reporter half-life. (**D**) Photoconversion results in a 40% drop in GFP intensity. As new Dendra2-LC3 is synthesized, GFP levels return to pre-photoconversion levels over the span of 13.5h. Torin1 blocks the observed return in GFP fluorescence by accelerating flux. Genetic inhibition of autophagy via siRNA-mediated knockdown of ATG5 2d prior attenuates Torin1’s effects in both the RFP (**C**) and GFP (**E**) channels. For plots B-D * indicates significant difference (p<0.05) using DMSO as reference group with Tukey’s multiple comparisons test. # indicates p<0.05 with the scramble control for each drug treatment as the reference group (i.e Scramble siRNA 1µM Torin1 vs. ATG5 siRNA 1µM Torin1). Superscript number indicates the first time point when significance was achieved.

To confirm autophagy-dependent degradation of Dendra2-LC3 in modified HEK cells, we asked whether genetic inhibition of autophagy extended Dendra-LC3 half-life. HEK cells were transfected with siRNA targeting the autophagy gene ATG5^49^, achieving a marked reduction in ATG5 levels (Fig. S1). ATG5 knockdown attenuated Torin1’s effects on Dendra2-LC3 half-life but had no discernible impact on Dendra2-LC3 turnover in vehicle treated cells (Fig. 2C). These data show that Dendra2-LC3 clearance in response to Torin1 is mediated by autophagy, and also suggest that ATG5 expression levels may be rate limiting only upon autophagy induction.

Consistent with effective photoconversion of Dendra2-LC3, GFP intensity dropped by approximately 40% in pulsed cells, returning to pre-conversion levels within 13h (Fig. 2D). This return to steady-state GFP intensity likely reflects an equilibrium point at which new Dendra-LC3 production is balanced with its turnover. Treatment with Torin1 shifted this balance, not only preventing the return in GFP signal, but also further reducing GFP intensity over time. Application of bafilomycin-A1 (Fig. 2D) or ATG5 knockdown (Fig. 2E) both prevented Torin1-induced reductions in Dendra2-LC3 GFP intensity. Conversely, bafilmoycin-A1 treatment led to a supra-physiological increase in GFP signal (Fig. 2D). Thus, while the decay of photoconverted (red) Dendra2-LC3 can be used to accurately measure autophagic flux because it decouples protein turnover and synthesis, time-dependent changes in native (green) Dendra2-LC3 fluorescence mirror those observed in the red channel, and provide complementary estimates of flux.

To confirm that the metabolism of endogenous Dendra2-LC3 reflects autophagic flux, while simultaneously validating the use Dendra2-LC3 cells for identifying new autophagy-modulating strategies, we used the cells to screen an Enzo tool compound library that includes several drugs with purported effects on autophagy (Fig. 3A, Figure 3-source data). These experiments helped gauge the generalizability of the assay beyond the effects of strong autophagy modulators such as Torin1 and bafilomycin-A1, and also helped determine its ability to identify drugs that impact autophagy through a variety of mechanisms.

**Figure 3:**
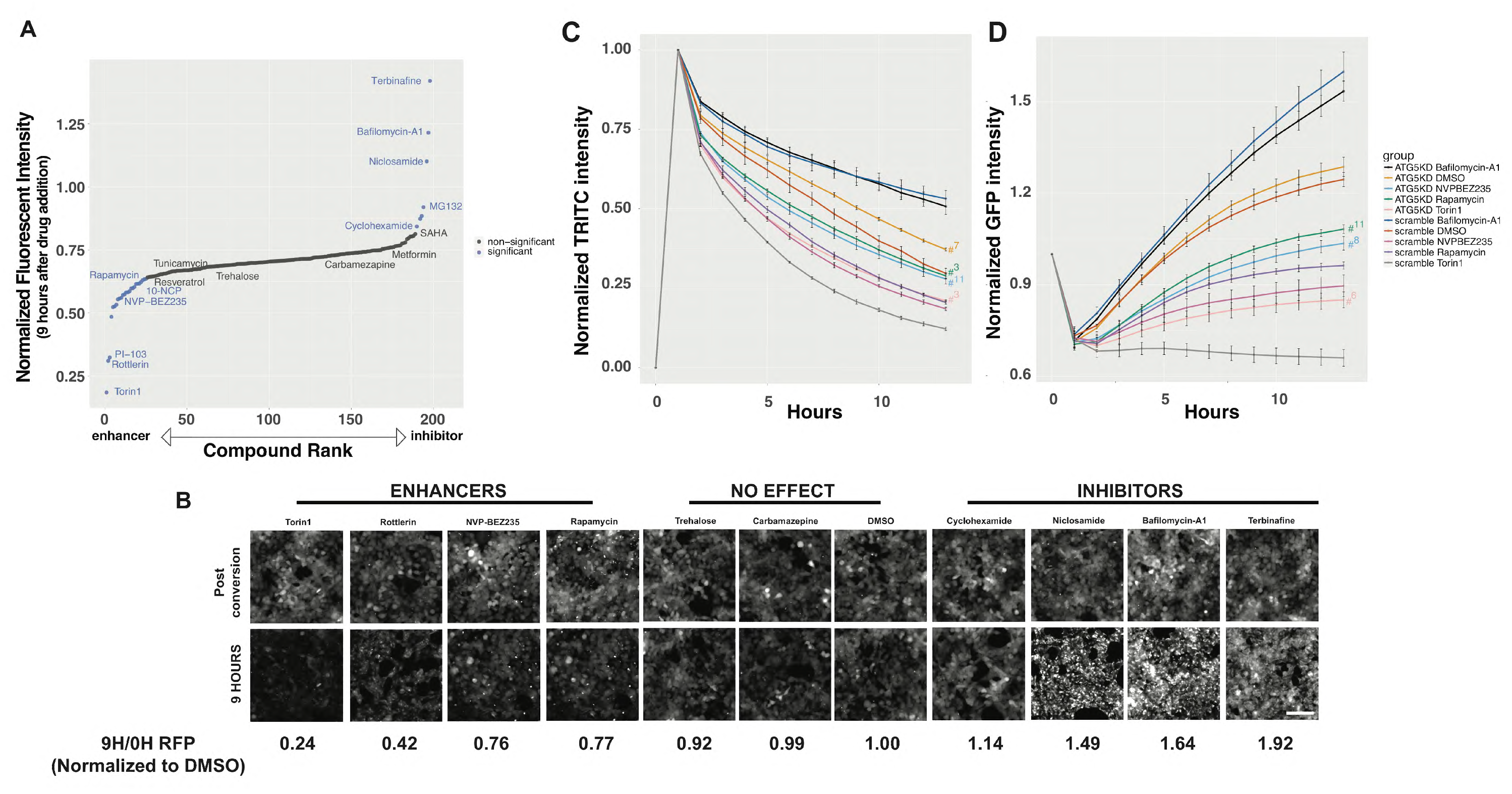
Small-scale screen in Dendra2-LC3 HEK293T cells confirm assay validity and identify new autophagy modulators. (**A**) An unbiased screen of the Enzo autophagy compound library identified several known autophagy-modulating compounds, including enhancers (rapamycin, NVP-BEZ235, AKT inhibitor X) and inhibitors (bafilomycin-A1). All drugs were added at a final concentration of 10µM via liquid handler, and autophagic flux estimated by their effects upon clearance of photoconverted (red) Dendra2-LC3 9h after drug addition. (**B**) Representative images of red Dendra2-LC3 immediately post-conversion and 9h after drug addition, demonstrating relatively rapid clearance with application of enhancers and marked accumulation of Dendra2-LC3 with inhibitors. Scale bar = 100µm (**C**) The effect of autophagy enhancers on Dendra2-LC3 half-life is attenuated by siRNA-mediated knockdown of ATG5 2d prior to drug application. Two-Way ANOVA indicated a significant interaction between time and treatment F(126,300)=15.22, p<0.0001 and significant effects for treatment F(9,300)=3427, p<0.0001 and time F(14,300)=957.3, p<0.0001 (**D**) Similar effects are observed in the green channel. Two-Way ANOVA found a significant interaction between time and treatment F(126,300)=20.71, p<0.0001. as well as significant main effects for treatment F(9,300)=721, p<0.0001 and time F(14,300)=320.8, p<0.0001. For (C) and (D) Error bars represent SEM from 3 replicate experiments. # indicates p<0.05 using Tukey’s multiple comparisons test with the scramble control for each drug treatment as the reference group (i.e Scramble siRNA Torin1 vs. ATG5 siRNA Torin1). Superscript number indicates the first time point when significance was achieved.

Cells were imaged once prior to photoconversion to measure background RFP intensity. As in previous experiments, a 4s pulse of 405nm light was used for photoconversion. Imaging occurred again immediately following photoconversion to determine the maximum RFP signal. All subsequent RFP intensities were normalized to this value after subtraction of the measured background RFP signal. Library drugs were added at 10µM via a robotic liquid handler, cells were imaged every 1.5h for 16h, and time-dependent changes in RFP intensity were measured for each condition. Using DMSO and Torin1 as negative and positive controls, respectively, we observed a Z’=.79 at 9h after drug application, demonstrating high sensitivity and reproducibility of the assay.

Many canonical autophagy-modulating drugs demonstrated clear effects on autophagic flux (Fig. 3A), establishing construct validity for the assay. Among compounds that significantly enhanced autophagic flux were the allosteric mTOR inhibitor rapamycin and the dual PI3K/mTOR inhibitors NVP-BEZ235 and PI-103. Torin1 and PI-103 exerted particularly strong effects in line with their action as ATP competitive antagonists^47, 50^. 10-NCP, an AKT inhibitor that we previously identified as a neuronal autophagy-inducing compound^40, 51^, also increased autophagic flux in Dendra2-LC3 HEK cells. Rottlerin, a compound that demonstrated autophagy-enhancing effects via inhibition of mTOR as well as protein kinase C (PKC) delta, greatly accelerated the degradation of Dendra2-LC3^52, 53^.

Bafilomycin-A1, also present in the compound library, registered as a strong inhibitor of Dendra2-LC3 flux (Fig. 3A), providing internal consistency with regards to our initial investigations. Rather than decrease over time, the intensity of photoconverted (red) Dendra2-LC3 in cells treated with bafilomycin-A1 progressively increased, eventually exceeding levels immediately following photoconversion (Fig. 3B). This phenomenon reflects a peculiar imaging property unique to Dendra2-linked proteins that accumulate within large puncta, in the process sequestering diffuse, low-intensity signal within relatively small regions (Fig. S2). Thus, the time-dependent degradation Dendra2-LC3 is inhibited by bafilomycin-A1, and the observed increase in red fluorescence intensity is due to the accumulation of Dendra2-LC3 within large clusters of perinuclear autophagosomes.

The protein translation inhibitor cyclohexamide stabilized Dendra2-LC3 turnover (Fig. 3A,B), in keeping with autophagy inhibition downstream of amino acid accumulation and mTORC1 activation^54^. This is in contrast to what is observed in the green channel, where inhibiting the synthesis of Dendra2-LC3 results in a decrease in GFP fluorescence as expected (Fig. 5-source data). These results highlight the pivotal ability of the assay to decouple autophagy inhibition from new protein synthesis.

Dendra2-LC3 RFP fluorescence was also stabilized by the proteasome inhibitor MG132 and the protease/proteasome inhibitor ALLN (Fig. 3A; Fig. 3-source data), albeit to a far lesser degree than with bafilomycin-A1 or other strong inhibitors. This suggests that while Dendra2-LC3 serves as valid reporter of autophagic flux, it is not degraded exclusively via autophagy. Therefore, to confirm that hits arising from this assay were indeed capable of affecting Dendra2-LC3 turnover via their actions on autophagy, we tested their effects in ATG5-deficient cells. As seen with Torin1 (Fig. 2), ATG5 knockdown attenuated the autophagy-inducing effects of NVP-BEZ235 and rapamycin (Fig. 3C,D), verifying that these drugs stimulate Dendra2-LC3 clearance by enhancing autophagic flux.

Because of the nature of the screen, compounds exhibiting intrinsic fluorescence could result in an artificially high RFP signal, leading to their subsequent misclassification as autophagy inhibitors. To address this possibility we rescreened all hits in unmodified HEK293 cells that do not express Dendra2. We found that 4 out of the 35 tested drugs, including 3 out of the 10 drugs that were identified as inhibitors, exhibited intrinsic fluorescence in the RFP channel (Fig. 3-source data, Fig. S3). For instance, curcumin, a purported autophagy modulator^55^ with known autofluorescent properties^56^ produced a substantial increase in background RFP signal that precluded any estimations of its effects on autophagy in this assay. In contrast, the PKC inhibitor bim-1 had little effect on background fluorescence, but instead accumulated within perinuclear autofluorescent puncta that resembled those observed in cells treated with bafilomycin-A1. These results underscore the importance of counter-screening to exclude intrinsically fluorescent drug properties that can confound or obscure results.

### Establishing a high-content screening platform for autophagy modulators

Using Torin1 and bafilomycin-A1, we next evaluated the sensitivity of Dendra2-LC3 cells for detecting small changes in autophagic flux in a dose-dependent manner. We tested the effects of 10 serial dilutions of each drug in both the GFP and RFP channels. As in previous experiments, baseline GFP and background RFP measurements were acquired prior to photoconversion. GFP measurements were normalized to this baseline value while RFP was normalized to the background-subtracted postconversion intensity. Following drug treatment, we imaged every 30m for 13.5h and anlayzed time- and concentration-dependent effects in each channel. We observed a tunable and proportional response to increasing drug concentrations for both Torin1 and bafilomycin-A1 (Fig. 4A). This was perhaps most evident for bafilomycin-A1, where the assay had sufficient resolution to discriminate 2nM changes in concentration (Fig. 4C, E). Notably, the GFP channel was nearly as sensitive as the RFP channel for detecting differences in autophagic flux (Fig. 4D, E). While the GFP fluorescence gradually returned to equilibrium in vehicle treated cells over a 12h span, it continued to drop with Torin1 treatment in a dose-dependent manner (Fig. 4D). Conversely, the GFP intensity quickly surpassed pre-photoconversion levels in cells treated with bafilomycin-A1, and the rate of increase was proportional to the drug dose (Fig. 4E). For Torin1, bafilomycin-A1, rapamycin and NVP-BEZ235, the dose response relationships for each drug were strikingly similar between channels, producing nearly identical half maximal effective concentration (EC50) and half maximal inhibitory concentration (IC50) values for each compound (Fig. 4F).

**Figure 4:**
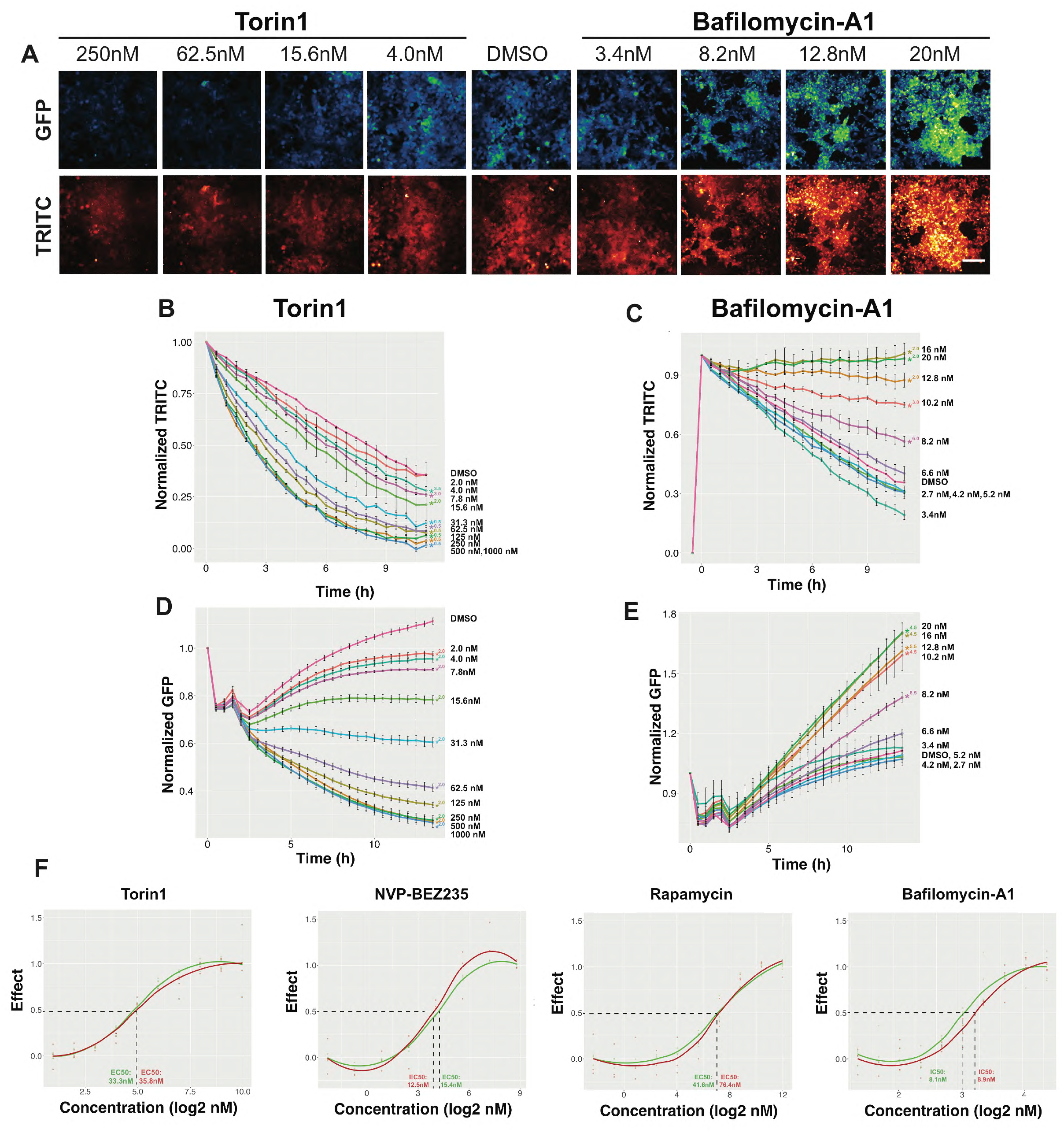
Proportional and bidirectional effects of autophagy modulators highlight assay sensitivity. (**A**) Dendra2-LC3 HEK293T cells were treated with increasing concentrations of Torin1 and bafilomycin-A1. Representative images in the GFP (top) and RFP (bottom) channels, pseudocolored to accentuate intensity variations. Scale bar = 100µm (**B**) Dendra2-LC3 clearance increased in a dose-dependent manner with Torin1 (**B**), while bafilomycin-A1 resulted in dose-dependent prolongation of Dendra2-LC3 half-life (**C**). These changes are even more apparent in the GFP channel for both Torin1 (**D**) and bafilomycin-A1 (**E**). Error bars represent SEM from 8 replicate wells. For plots B-E * indicates significant difference (p<0.05) using DMSO as reference group with Dunnett’s multiple comparisons test. Superscript number indicates the first time point when significance was achieved. (**F**) Dose-response curves for Torin1, NVP-BEZ235, rapamycin, and bafilomycin-A1. For autophagy enhancers, the minimal RFP intensity 7h after drug treatment relative to DMSO was set to 1, and the maximal value set to 0. For inhibitors, the maximum effect represents the maximal RFP intensity within 7h after drug treatment. Dose-response was determined similarly for the GFP channel, utilizing values 14h after drug treatment. Concentration is plotted in nM on a log(2) scale, with ≥ 3 replicate wells for each channel shown as colored dots. EC50 and IC50 values are reported along the x-axis for both RFP and GFP.

These data indicate that both the GFP and RFP channels provide accurate information regarding autophagic flux upon drug addition. Since imaging in the GFP channel does not require photoconversion, experiments take only a fraction of the time that would otherwise be needed to track Dendra-LC3 turnover in the RFP channel. We took advantage of this fact in developing a high-throughput and high-content screening platform in Dendra2-LC3 HEK cells. Cells were imaged in the GFP channel immediately prior to drug addition (GFP_0H_) and then again 15h later (GFP_15H_). Autophagy enhancers were defined as those drugs that reduce the GFP_15H_/GFP_0H_ ratio by more than 3 standard deviations from the mean of vehicle (DMSO), and inhibitors as drugs that increase the GFP_15H_/GFP_0H_ ratio by more than 3 standard deviations. In 3 replicate 90/10 studies where 90% of wells were treated with DMSO and 10% were treated with Torin1, we observed a 1% false positive rate, no false negatives, and a mean Z’=.52, thus validating this method as a reliable primary screening assay (Fig. 5A, Table 1).

**Figure 5:**
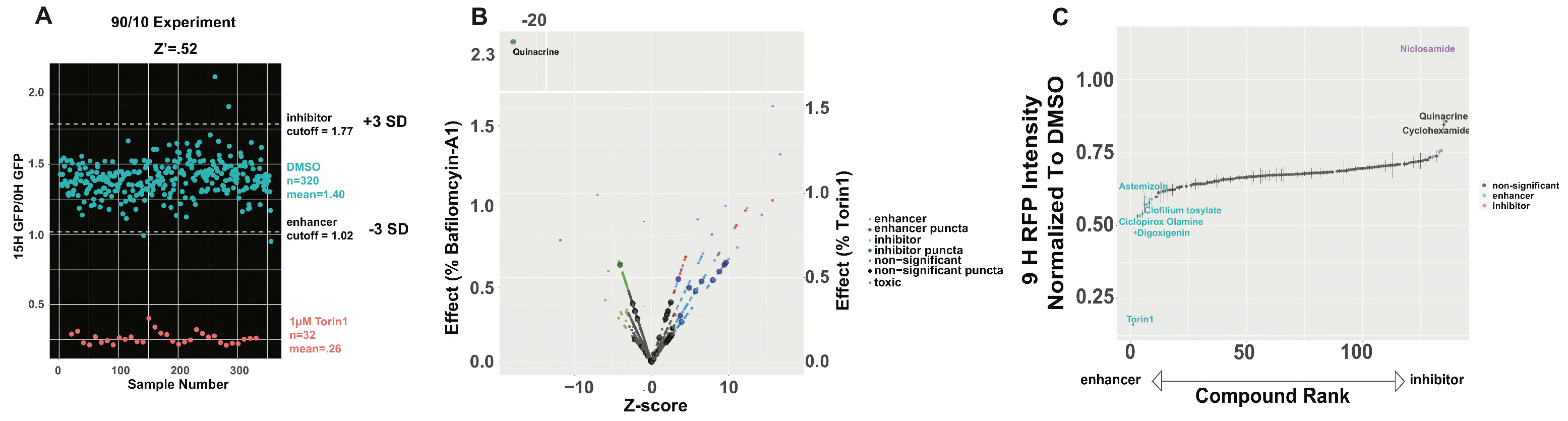
Screening the Prestwick library reveals previously unrecognized autophagy-modulating drugs. (**A**) “90/10” experiment validating the measurement of Dendra-LC3 in the GFP channels as a primary assay. 320 wells of a 384w plate were treated with DMSO and 32 were treated with 1µM Torin1. Plates were imaged in GFP immediately before, and 15h after drug treatment. Z’=.52 ± 0.04 in 3 replicates. (**B**) Primary screen of the Prestwick drug library. Z-score is calculated as number of SD_DMSO_ greater or less than mean DMSO_15H/0H_. Significant changes in flux, toxicity, or puncta for each drug are indicated by the color and size of representative dots, according to the key. The magnitude of effect is represented as both % bafilomycin-A1 (left y-axis) and % Torin1 (right y-axis) (**C**) Time-dependent decay in red (photoconverted) Dendra2-LC3 was used as a secondary screen of the non-toxic candidates emerging from the primary screen. Error bars represent the SEM from 6 images (2 images/well of 3 replicate wells).

**TABLE 1:** 90/10 experiments evaluating primary high-throughput screening assay

|  | Experiment 1(n=96) | Experiment 2 | Experiment 3 |
| --- | --- | --- | --- |
| Z' | 0.51 | 0.49 | 0.56 |
| false positives | 1 | 3 | 4 |
| false negatives | 0 | 0 | 0 |
| mean DMSO-mean Torin1 | 0.75 | 1.46 | 1.14 |
| DMSO SD/DMSO mean | 6.40% | 11% | 9% |

We then devised a layered screening scheme where hits from the primary assay were filtered based on toxicity, then subjected to a counter-screen where Dendra2-LC3 half-life is determined following photoconversion and imaging in the red channel. This organization combines the added throughput of imaging in the green channel with the ability to selectively monitor Dendra2-LC3 degradation in the red channel. Custom scripts were used to exclude toxic compounds based on their effects on cell number— this was particularly important since drugs that cause cells to die might significantly lower GFP intensity and could therefore be misconstrued as false-positives.

We first applied this screening strategy to the Prestwick drug library, a collection of 1,280 off-patent small molecules, 95% of which have gained regulatory approval by the FDA, EMA and other agencies. Eighteen compounds were filtered out due to toxicity; 17 of which would have otherwise been identified as autophagy enhancers due to their ability to significantly reduce GFP intensity. Among the remaining compounds, 129 exhibited significant effects on Dendra2-LC3 levels, with 88 significantly reducing the GFP_15H_/GFP_0H_ ratio (i.e. enhancing autophagy) and 41 increasing the ratio (i.e. inhibiting autophagy) (Fig. 5B, Fig. 5-source data).

We also assessed the frequency of Dendra2-LC3 puncta—corresponding to autophagosomes—in treated cells, since a change in the number or size of puncta could indicate either autophagy induction or late-stage autophagy inhibition^3^. In fact, 2 previous high-throughput screens utilized changes in LC3 puncta number to identify autophagy modulators^57, 58^. In native Dendra2-LC3 cells, we observed an increase in Dendra2-LC3 puncta number in response to 38 drugs, but only 13 of these compounds significantly affected autophagic flux in the primary screen. Among these 13, 11 reduced the GFP_15H_/GFP_0H_ ratio and were counted as enhancers, while 2 compounds acted as inhibitors and increased the GFP_15H_/GFP_0H_ ratio (Fig. 5B, Fig. 5-source data).

All 129 hits, along with 2 drugs that narrowly missed significance in the primary assay, were then evaluated for their ability to affect the degradation of photoconverted Dendra2-LC3. 17 compounds (13%) significantly modulated autophagic flux; of these, 11 enhanced flux and 6 inhibited it. Counter-screening in unmodified HEK cells uncovered two drugs, diacerin and mitoxanthrone, with intrinsic red fluorescence independent of their effects on autophagy Fig. 5-source data, Fig. S3).

Among notable enhancers were the antiarrhythmic drugs digoxigenin, a metabolite of digoxin, and clofilium tosylate. Ciclopirox olamine, an off-patent anti-fungal agent found in both the Prestwick and Enzo libraries, potently enhanced autophagy. Similarly, the anthelmintic niclosamide inhibited autophagy to a comparable extent as bafilomycin-A1 in the Enzo compound screen, and was the strongest inhibitor tested in the Prestwick library, demonstrating significant consistency of the assay across different libraries. As seen previously, cyclohexamide modestly stabilized photoconverted (red) Dendra2-LC3, despite lowering green Dendra2-LC3 levels in the primary screen. This ability to exclude compounds that lower LC3 levels due to translation inhibition demonstrates the value of screening in both the GFP and RFP channels. These data, along with the observation that 87% of primary screen “enhancers” failed to stimulate autophagy in counter-screening, suggest that the primary screen is a sensitive but not specific method to identify autophagy enhancers, but the added selectively of the secondary screen provides a means to successfully filter out false enhancers.

### An expanded screen identifies novel autophagy inhibitors

Our data indicate that Dendra2-LC3 cells provide a robust, accurate and precise measure of autophagic flux that can be readily adapted for compound screening. To identify new autophagy modulating drugs, we took advantage of the screening assay we developed to investigate a library of 24,000 drugs spanning considerable chemical diversity curated from the Maybridge library. In our screen of the Prestwick library a large percentage of hits from the primary screen (time-dependent changes in the GFP intensity) failed to show an effect in the secondary assay (time-dependent changes in RFP intensity post-photoconversion). Since we aimed to test ∼20 times as many compounds as before, we chose to filter out false positives by repeating the primary assay and retesting hits after exclusion of toxic compounds, before progressing to secondary screening involving clearance of photoconverted Dendra2-LC3. We also confirmed each hit by repeating the secondary screen prior to further filtering based on the solubility and permeability of each hit. Finally, the intrinsic fluorescence of each candidate compound was assessed in unmodified HEK239T cells, and all remaining hits were confirmed using orthogonal flux assays (Fig. 6A).

**Figure 6:**
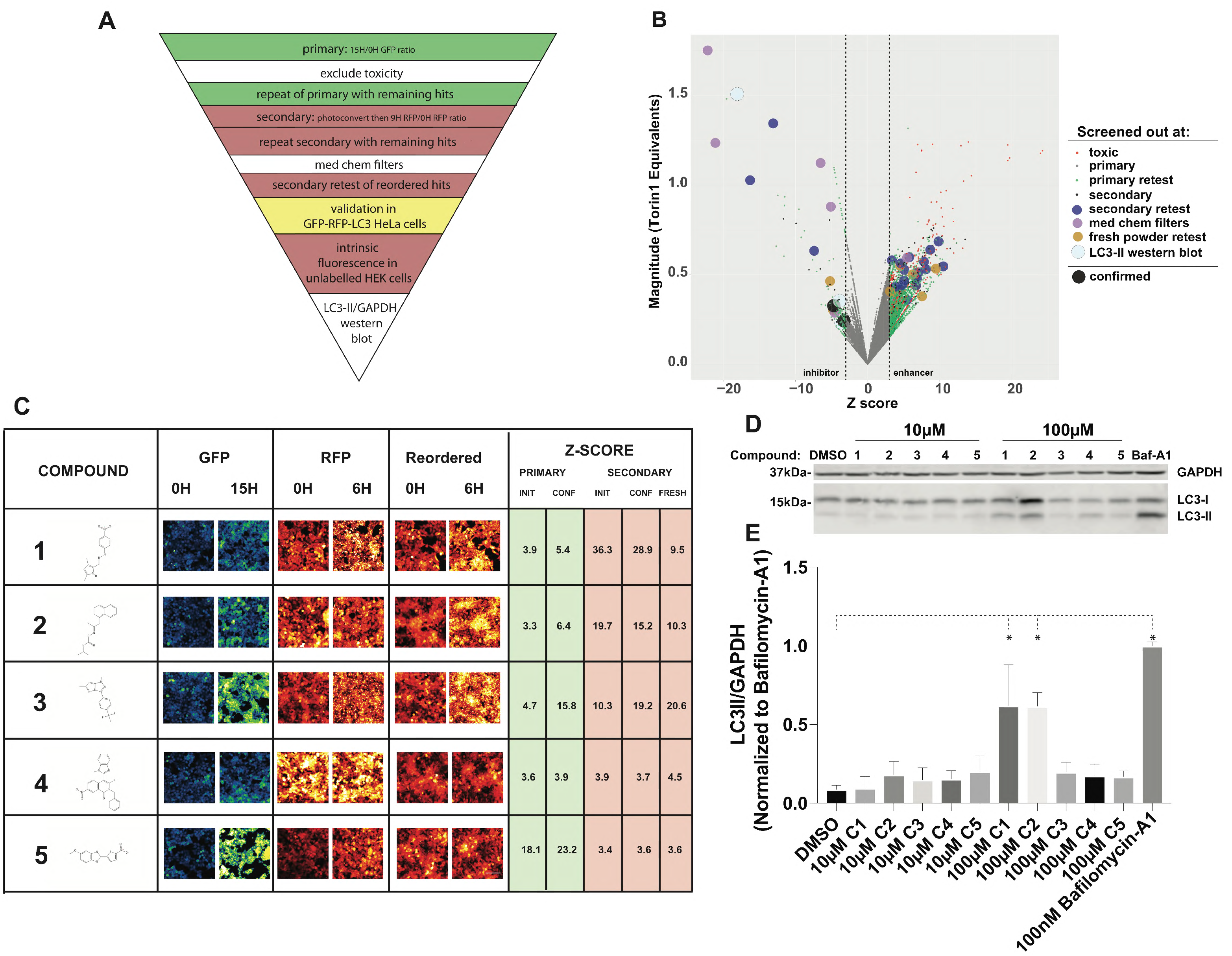
High throughput screening identifies novel autophagy inhibitors. **A)** Schematic depicting the screening hierarchy used. **(B)** The primary screen was performed using the Maybridge 24K library, consisting of 24,000 chemically diverse compounds. For enhancers, changes in Dendra2-LC3 GFP intensity were normalized to Torin1’s effects. Non-toxic compounds that passed the primary screen were filtered by re-testing in the GFP channel, then evaluated in a secondary screen involving calculation of Dendra2-LC3 half-life in the RFP channel. Hits were re-tested in the secondary screen, followed by repeat evaluation in the RFP channel using fresh compound from a different distributor. The color and size of each dot denotes the stage at which individual compounds were eliminated, in accordance with the key. Candidates that passed all filters are shown in black. (**C**) Representative images of the primary screen, secondary screen, and repeat secondary screen with fresh drug for the 5 putative autophagy inhibitors, ranked based on the magnitude of inhibition measured in the initial secondary screen. Z-scores are reported for each screening phase. Scale bar = 100µm. (**D**) Unmodified HEK293T cells were treated with vehicle or each compound at either 10µM or 100µM. 9H after treatment lysates were collected and immunoblotted with an LC3 antibody. (**E**) Quantification of three replicate experiments demonstrating that at 100µM of compounds 1 and 2 significantly inhibit autophagic flux. For each group LC3-II was normalized to the loading control GAPDH and then scaled to 100nM bafilomycin-A1. Error bars represent standard error of the mean. One-way ANOVA showed significant differences between groups (F=24.28, P<0.0001). * p<0.01 compared to DMSO, Dunnet’s multiple comparison test.

Similar to the 10% hit rate observed with the Prestwick library, we identified 2160 compounds (9%) as potential autophagy modulators from the primary screen. Of these, 1958 registered as autophagy enhancers, and 202 as inhibitors. Upon repeating the screen after excluding toxic compounds, 232 candidates (1%) remained as hits, underscoring the need to repeat the primary screen. The retest reduced the number of enhancers more than 10-fold while decreasing the number of inhibitors by a factor of 2.4, demonstrating a predilection towards false-enhancers in the primary screen. Following the secondary screen and retest, 23 compounds remained. Notably, all enhancers identified in the primary screen failed to pass the secondary screens (Fig, 6B, Table 2).

**TABLE 2:** Summary of Maybridge library screen and orthogonal autophagic flux assays.

| Phase | # compounds | % | Enhancer | Inhibitor |
| --- | --- | --- | --- | --- |
| Maybridge Library | 24000 | 100 | NA | NA |
| Primary | 2160 | 9 | 1958 | 202 |
| Primary confirmation | 232 | 0.97 | 148 | 84 |
| Secondary | 41 | 0.17 | 3 | 39 |
| Secondary confirmation | 23 | 0.1 | 0 | 23 |
| passed med chem filters | 16 | 0.07 | 0 | 16 |
| retest reordered drugs | 5 | 0.02 | 0 | 5 |
| GFP-RFP-LC3 HeLa | 5 | 0.02 | 0 | 5 |
| LC3II western blot | 2 | 0.01 | 0 | 2 |

Of the 23 candidate autophagy inhibitors, 7 were excluded based on limited solubility and permeability, and 11 more were discarded upon acquisition of fresh powder from commercial sources. Among the 5 remaining autophagy inhibitors (Fig 6C, Table 2), the top 3 candidates consistently tested among the most potent inhibitors in all assays. Additionally, each compound elicited a dramatic perinuclear accumulation of Dendra2-LC3 in modified HEK293T cells (Fig. 6C), suggestive of a late-stage block in autophagosome maturation.

We next sought to validate the ability of these compounds to inhibit autophagy using an alternative flux assay. To do so we utilized a HeLa cell line stably overexpressing the tandem RFP-GFP-LC3 reporter^59^. In this system, LC3 is fused to an acid-sensitive GFP as well as an acid-insensitive RFP. Upon progression from autophagosome to autolysosome, GFP fluorescence is quenched as the sensor enters an acidic environment. Application of autophagy inhibitors such as bafilomycin-A1 that inhibit lysosomal acidification result in the appearance of GFP(+)/RFP(+) (yellow) autophagosomes (Fig. S4A). All 5 of the newly-identified compounds significantly increased yellow puncta accumulation in RFP-GFP-LC3 HeLa cells, indicative of effective autophagy inhibition. To assess this effect in an automated and unbiased manner, we developed an image analysis pipeline that identifies cytoplasmic puncta and reports the fraction of GFP(+)/RFP(+) puncta (Fig. S4B). Using this pipeline we observed dose-dependent effects for each compound across similar concentration ranges as those observed in Dendra2-LC3 HEK293T cells (Fig. S4C; Fig. S5). After 12h, all compounds save #2 had reached their maximal response (Fig. S4D). Compounds 1,4, and 5 exerted more than half their maximum effect immediately after drug addition. Quinacrine, a previously reported autophagy inhibitor^60^, showed similar kinetics (Fig.S4E), while the response of compound 3 more closely matched the kinetics of bafilomycin-A1.

We next asked whether these compounds exhibited intrinsic fluorescence that might interfere with our assessment of autophagic flux. Compound 4 did not produce red fluorescence at any concentration tested, but we observed dose-dependent red fluorescence for all other compounds (Fig. S5A). Even so, each compound inhibited autophagy in both flux assays at concentrations where intrinsic fluorescence is minimal (Fig. S5B). In the green channel, all drugs produced significant green fluorescence with the exception of compound 4. Application of compound 3 resulted in the appearance of bright perinuclear puncta that could confound estimates of GFP(+)/RFP(+) puncta in HeLa cells. Indeed, in this assay a lower concentration of compound 3 was needed to exert an effect than in the Dendra2-LC3 flux assay, which should be unaffected by changes in green fluorescence (Fig. S5B).

To exclude any possible contribution of fluorescence in the measurement of flux, we assessed autophagy inhibition using an orthogonal assay that does not rely on fluorescent readouts. Treatment with compounds 1 and 2 resulted in a significant and reproducible accumulation of LC3-II by western blotting, indicative of impaired autophagosome clearance (Fig. 6D,E). Together these data demonstrate that both compounds are indeed autophagy inhibitors; however, the intrinsic fluorescence of compound 2, and to a lesser extent compound 1, can complicate readouts of flux in fluorescence-based autophagy assays.

### Enhancing autophagic flux suppresses toxicity in a primary neuron model of ALS

We previously demonstrated that autophagy induction using a family of small molecules protected against TDP43-mediated toxicity in models of ALS/FTD^40^. Here we sought to test whether any enhancers identified in our compound screens conferred similar neuroprotection. To accomplish this we turned to automated fluorescence microscopy (AFM), a technology capable of individually tracking thousands of cells prospectively over time^61, 62^. Primary rodent spinal and cortical neurons were transfected with Dendra2-LC3, photoconverted with a brief pulse of blue light and imaged by AFM (Fig. 7A). Because the platform measures single-cell intensity within the TRITC channel over time, we are able to calculate a half-life for Dendra2-LC3, corresponding to autophagic flux, within each neuron. Cortical neurons exhibited slightly higher basal rates of autophagy than spinal neurons, with a mean single-cell Dendra2-LC3 half-life of 33.2h compared to 37.1h seen in spinal neurons (Fig. S6A,B, p=7.1×10^-4^, Welch two sample t-test).

**Figure 7:**
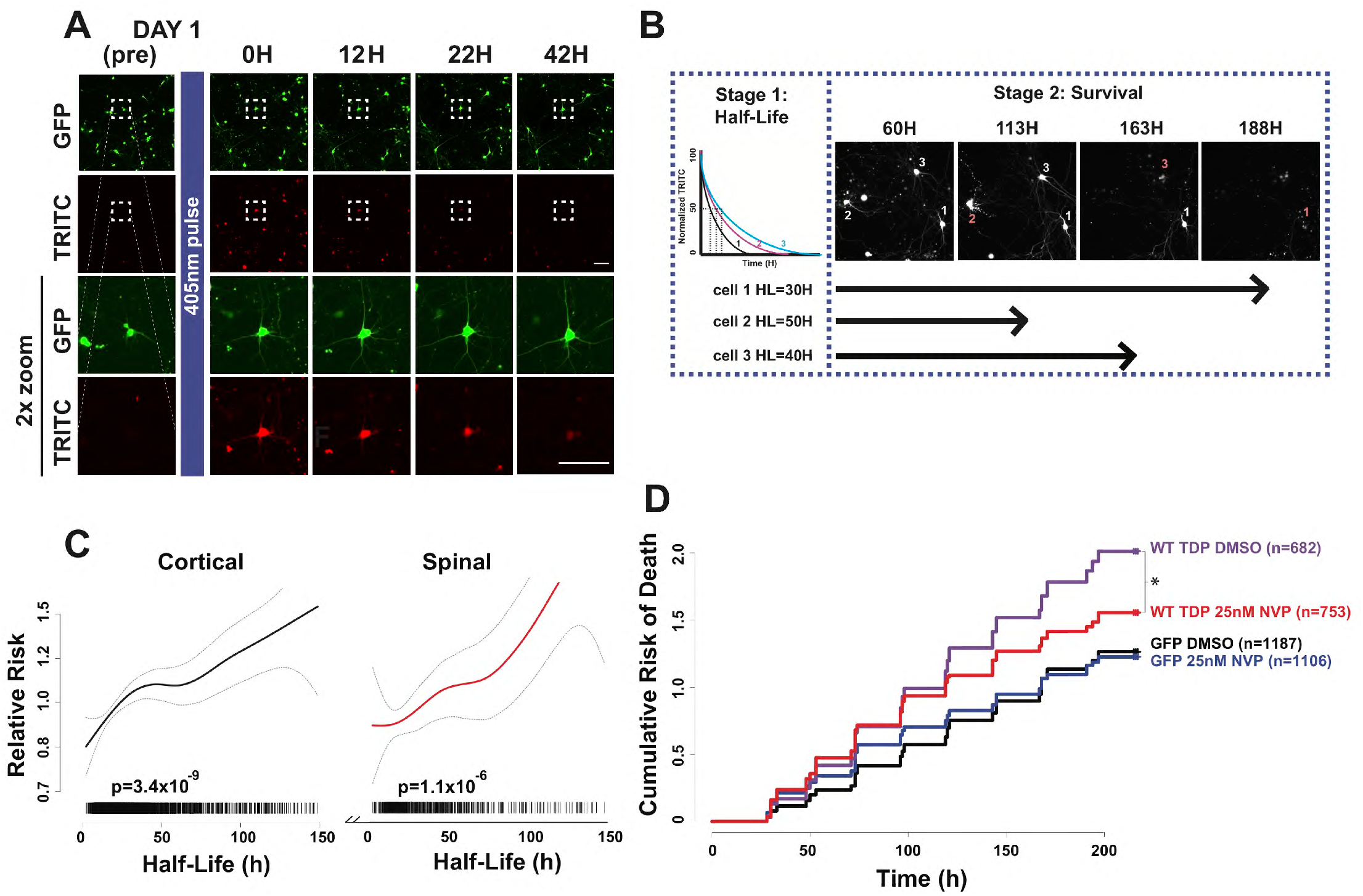
Autophagic flux predicts neuronal survival and mitigates toxicity in an ALS/FTD disease model. (**A**) Mixed rat spinal neurons were transfected on DIV 4 with Dendra2-LC3, imaged 24h later (DAY 1 pre), then pulsed with 405nm light to photoconvert Dendra2-LC3 before imaging repeatedly and longitudinally over several days to track the time-dependent loss of red fluorescence and neuronal survival. Scale bars = 100µm in each panel. (**B**) Experimental outline for determining the relationship between Dendra2-LC3 half-life and neuronal survival. After calculating Dendra2-LC3 half-life for individual neurons (Stage 1), each cell is prospectively tracked using automated fluorescence microscopy to determine its time of death (Stage 2; red number and corresponding arrow). (**C**) Penalized spline Cox proportional hazards model depicting Dendra2-LC3 half-life (x-axis) vs. relative risk of death (y-axis) for primary rat cortical (black) and spinal (red) neurons, demonstrating a strong linear relationship for both populations (cortical: p=3.4×10^-9^; spinal p=1.1×10^-6^, linear Cox proportional hazards). Cortical and spinal neurons with shorter Dendra2-LC3 half-lives, and therefore higher rates of basal autophagy, lived longer. Each hash mark represents an individual neuron, collected from 3 biological and 8 technical replicates each. Grey dotted lines mark 95% confidence intervals. (**D**) NVP-BEZ235 (25nM NVP) treatment suppresses toxicity in primary rat cortical neurons overexpressing WT-TDP43-GFP. Table 3 summarizes the hazard ratio and statistical significance of each comparison as determined by Cox proportional hazards analysis. N for each group represents total neurons pooled from three replicate experiments. * p<0.05, cox proportional hazards analysis.

**TABLE 3:** Summary of Cox proportional hazards analysis

| Comparison | Hazard Ratio | p-value |
| --- | --- | --- |
| GFP/DMSO vs. TDP43-GFP/DMSO | **1.74** | **1.02E-18** |
| GFP/DMSO vs. GFP/25nM NVP-BEZ235 | 1.02 | 0.67 |
| WT-TDP43-GFP DMSO vs. WT-TDP43-GFP/<br>25nM NVPBEZ235 | **0.89** | **0.03** |

We also determined the lifespan of each neuron using custom scripts that assign a time of death for each cell^62, 63^. To assess the relationship between basal rates of autophagy in neurons and their survival, we incorporated Dendra2-LC3 half-life as a continuous variable into a Cox proportional hazards model of neuronal survival^64^ (Fig. 7B). For both cortical and spinal neurons, rapid turnover of Dendra2-LC3 was associated with extended lifespan (Fig. 7C, cortical: p=3.4×10^-9^, spinal p=1.1×10^-6^, Cox hazards analysis). These results indicate that higher rates of basal autophagic flux are associated with prolonged neuronal survival *in vitro*, providing further rationale for the development of autophagy activators for neurodegenerative diseases.

NVP-BEZ235 (dactolisib) was among the most potent autophagy enhancers identified using Dendra2-LC3 HEK cells (Figs. 3,4). In light of our prior data attesting to the benefit of autophagy induction in ALS/FTD models and the proportional relationship between autophagic flux and neuronal survival (Fig. 7C), we predicted that NVP-BEZ235 could prevent neurodegeneration in disease models. We therefore tested this compound in neurons overexpressing TDP43, an RNA binding protein whose accumulation is integrally connected with both ALS and FTD^65, 66^. In prior studies, TDP43 overexpression reproduced characteristic features of disease, including the formation of ubiquitin- and TDP43-positive neuronal inclusions, cytoplasmic TDP43 mislocalization, and neurodegeneration^40, 67, 68^. As expected, TDP43 overexpression resulted in an increase in the risk of death in comparison to neurons transfected with EGFP, a control protein. Treatment with NVP-BEZ235 significantly prolonged the survival of neurons overexpressing TDP43 (p<0.05, HR=0.89, Cox hazards analysis), without adversely affecting EGFP-expressing neurons (Fig. 7D, Table 3). These data demonstrate the therapeutic potential of NVP-BEZ235 as an autophagy inducer capable of extending neuronal survival in ALS/FTD models.

## Discussion

In this study we developed a unique human reporter cell line enabling the non-invasive measure of autophagic flux in living cells, without interference from pathway intermediates or potentially toxic pathway inhibitors. Building on previous efforts to isolate autophagic clearance from induction^40, 59, 69^, here we created a system with the pivotal capacity to assess *native* autophagic flux, thereby avoiding several basic confounds associated with overexpression of autophagy reporters^70, 71^. This is of particular relevance considering the intersection between autophagy and nutrient/energy sensing^42^, the role of microtubule associated transport in autophagosome maturation^48^, and the crosstalk between autophagy and the ubiquitin proteasome system^44^. Increasing protein dosage can also induce aberrant aggregation of misfolded proteins, and influence the likelihood of protein phase separation^70, 72^. Beyond the potential toxicity associated with these outcomes, overexpression of LC3 and LC3-based reporters is sufficient to produce visible puncta that could be mistaken for autophagosomes.

Through the targeted insertion of Dendra2 into the *MAP1LC3B* locus, we generated reporter cell lines in which Dendra2-LC3 expression is driven by the endogenous *MAP1LC3B* promoter. As such, the baseline fluorescence intensity of non-photoconverted (green) Dendra2-LC3 reflects the steady-state balance between LC3 synthesis and degradation. We took advantage of this relationship to quickly and accurately gauge the effects of compounds that enhance or inhibit autophagy, without the need for photoconversion of Dendra2-LC3. Such an approach can be problematic in overexpression-based systems, but it provided a robust means of estimating autophagic flux on a high-throughput basis in Dendra2-LC3 HEK cells. Future studies employing a full bleach or photoconversion of Dendra2-LC3 could be useful for investigating autophagy regulation at the level of transcription or protein synthesis, and for identifying genetic and/or pharmacologic approaches that induce autophagy at an early stage.

Previous high-throughput screens for autophagy modulators utilized the formation of LC3-positive puncta as the major criterion for autophagy induction^57, 58^. As late stage autophagy inhibition is equally effective as autophagy induction for triggering the accumulation of LC3 puncta, such analyses are unable to discriminate whether a drug truly enhances or blocks flux. For example, the drugs loperamide and astemizole, two drugs found here to enhance flux, were also identified as potential autophagy inducers based on their ability to increase LC3 puncta^58^. However, niclosamide was also labeled as an inducer because of its effect on LC3 puncta, but was in fact a strong <u>inhibitor</u> of flux in Dendra2-LC3 cells. Further supporting the apparent disconnect between puncta number and autophagic activity, we identified several compounds that increased Dendra2-LC3 puncta without markedly impacting Dendra2-LC3 levels (Fig. 5A). As such, reporters that judge the clearance of autophagy substrates are intrinsically more suited to gauging autophagic flux than are those that focus on pathway intermediates such as autophagosome number. Because autophagy is a dynamic pathway that requires coordinated regulation of several critical steps, increasing autophagy *induction* without an accompanying downstream increase in substrate clearance is likely to be of little therapeutic benefit and may even be maladaptive^73, 74^. Therefore, we placed particular emphasis on enhancing *productive* autophagy and the measurement of autophagic flux through the use of non-invasive reporters.

In comparison to alternative methods for measuring autophagic flux, Dendra2-LC3 cells offer unique advantages. Not only do these cells afford the only means of estimating *native* autophagic flux in living cells, but they also preclude the need for overexpression of LC3 analogues, thereby avoiding many of the pitfalls that plague other approaches. For instance, while the GFP-LC3-RFP-LC3ΔG reporter^69^ is likewise capable of discriminating LC3 synthesis from degradation, measurements of autophagic flux using this probe require relating the degradation rates of two proteins, one of which is an autophagy marker and substrate (GFP-LC3) and another that is untethered within the cytoplasm (RFP-LC3ΔG). Conditions that stabilize RFP-LC3ΔG without accelerating GFP-LC3 clearance could be misinterpreted as increasing flux. Supporting this notion, in a screen using the GFP-LC3-RFP-LC3ΔG reporter, MG132 was identified as an autophagy enhancer^69^, while we found that MG132 instead stabilized Dendra2-LC3. Conversely, compounds such as loperamide that enhance proteolytic clearance via the ubiquitin-proteasome pathway^75^ may be mislabeled as autophagy inhibitors because of their preferential effects on RFP-LC3ΔG.

Several drugs purported to stimulate autophagy—including trehalose^76, 77^, metformin^78, 79 80, 81^, and carbamezapine^82, 83, 84^—failed to do so in our Dendra2-LC3 cell line. Despite promising results in mouse models of Huntington’s disease^76^, ALS^85^, and Parkinson’s disease^86^, evidence that trehalose induces autophagy is variable, with some studies claiming that the drug actually inhibits flux^87, 88, 69^. Likewise, the ability of carbamazepine to enhance autophagic flux and prevent neurodegeneration was based upon changes in steady-state levels of autophagy intermediates^83, 84^. Our data suggest that the protective effects of these drugs may not be a result of autophagy stimulation. However, the discrepancy in findings may be due to the high concentration of these drugs used in previous studies^81, 79, 80, 89, 77, 82^ relative to the 10uM dose used in our screens, as well as species and cell type specific differences.

Our screen of the Maybridge library suggests that uncovering novel autophagy enhancers may be considerably more challenging than inhibitors. Testing larger libraries or incorporating iterative chemical synthesis guided by structure-activity relationships^50^ may be required to effectively identify new autophagy-inducing compounds. Even so, compounds 1 and 2 were repeatedly found to inhibit autophagy in Dendra2-LC3 cells, RFP-GFP-HeLa cells, and via immunoblotting. While these drugs hold potential for the treatment of neoplasms that rely on autophagy for survival, their potency, activity, and bioavailability could be improved through similar means.

Despite the broad therapeutic potential of autophagy modulation, there are no clinically-approved drugs that have been specifically developed to target autophagy^7^. Our work with NVP-BEZ235 presents a potential template to evaluate the efficacy of putative autophagy modulators. We validated the autophagy-enhancing effects of the compound in Dendra2-LC3 HEK293T cells, and further utilized these cells to establish a dose-response relationship that informed subsequent studies demonstrating the drug’s therapeutic effects in a model of ALS and FTD. Given previous data indicating the ability of NVP-BEZ235 to suppress amyloid-β induced neurotoxicity^90^, and our results showing its potential in a model of ALS and FTD, this compound may be broadly effective in stimulating autophagy and preventing neuron loss arising from the accumulation of misfolded proteins in Alzheimer’s disease as well as ALS, FTD, and related conditions.

Autophagy has alternatively been reported to play protective or detrimental roles in ALS^91, 92^. In our previous work, pharmacological activation of neuronal autophagy suppressed TDP43 mediated toxicity^40^. This is in accordance with the protective effects of rapamycin in a TDP43 mouse model^84^. In contrast, rapamycin administration in SOD1 mice exacerbated muscle degeneration and shortened lifespan^93^. One study in which autophagy was genetically ablated from motor neurons in SOD1 (G93A) mice provided further insight into the conflicting roles of autophagy in ALS^73^. Loss of autophagy in motor neurons accelerated disease onset but also prolonged survival. This study suggested that early in disease autophagy plays a critical role in the maintenance of neuromuscular junctions. However, at later stages it promotes disease progression in a non-cell-autonomous manner. Our work further supports a protective role of autophagy enhancement in ameliorating toxicity associated with the accumulation of TDP43. Future studies may determine whether NVP-BEZ235 confers similar beneficial effects in other disease models involving SOD1 or RNA binding proteins related to TDP43. Even so, neurons respond poorly to most autophagy-inducing stimuli, making them a particularly challenging cell type to target for therapy development^51, 94, 95^. Additional investigations involving the introduction of Dendra2 into the *MAP1LC3B* locus of induced pluripotent stem cells may be critical for defining the cell type-specific mechanisms governing autophagy within neurons and other cell types.

While this technology represents a considerable advancement, it is not without limitations. Chief among these is the reliance on reductions in Dendra2-LC3 signal intensity to indicate enhanced flux. For this reason, we developed automated programs to selectively remove drugs with toxic effects that might otherwise be misclassified as false-positives. Since HEK293T cells are actively dividing, compounds that merely inhibit growth rate may also be falsely identified as enhancers when measuring GFP fluorescence, necessitating the use of counter-screens involving the measurement of photoconverted (red) Dendra2-LC3 half-life to eliminate these false-positives from the final pool of candidate compounds. We imaged multiple frames/well and multiple wells to account for autofluorescence artifacts that are common in the red channel, but the use of brighter fluorophores or photoconvertible fluorescent proteins that emit in the far-red wavelengths may avoid these complications^96^.

Additionally, due to the assay’s reliance on measuring fluorescent intensity, drugs that exhibit intrinsic fluorescence can confound flux measurements. In unlabeled HEK293T cells, compound 3 from the Maybridge library screen and the drug Bim-1 accumulated within bright fluorescent perinuclear puncta with striking resemblance to puncta observed in bafilomycin-A1 treated Dendra2-LC3 HEK293T cells (Fig. S3). We therefore counter-screened all candidates in unlabeled HEK293T cells and eliminated those exhibiting intrinsic fluorescence at the concentrations tested. We also confirmed autophagy inhibition using a non-imaging based flux assay. Despite the fact that all 5 Maybridge library hits demonstrated autophagy-inhibiting properties in RFP-GFP-LC3 expressing HeLa cells, immunoblotting revealed that only compounds 1 and 2 produced a significant increase in LC3-II levels. Therefore, due to its reliance on measuring fluorescence intensity, the RFP-GFP-LC3 flux assay suffers from the same problem in misidentifying intrinsically fluorescent drugs as inhibitors. Any fluorescence-based autophagy assays are likely to be similarly impacted by intrinsic fluorescence, emphasizing the need to account for these effects in screening efforts.

Ideally, future studies will evaluate autophagic flux using complimentary reporters that provide valuable information on mechanism of action, in addition to magnitude and potency. Such an approach, coupled with a shift toward analyzing the productive autophagic clearance of substrates expressed at endogenous levels, promises to accelerate and facilitate the discovery of novel autophagy-modulating compounds with wide-ranging therapeutic potential.

## Methods

### Cell Culture

HEK293T Dendra2-LC3 and HeLa RFP-GFP-LC3 cells were cultured in HEK complete media consisting of Dulbecco’s Modified Eagle Medium (DMEM) (GIBCO, Waltham, MA, cat #: 11995-065) supplemented with 20% fetal bovine serum, 1% GlutaMAX (GIBCO, cat #: 35050-061) and penicillin-streptomycin. For imaging experiments cells were placed in Neumo media (Cell Guidance Systems, Cambridge, UK, cat #: M07-500) with SOS supplement (Cell Guidance Systems, cat #: M09-50).

### CRISPR editing

Single guide RNA pairs (sgRNAs; Table 4) were selected using the CRISPR design tool available at http://crispr.mit.edu/. Sense and antisense oligonucleotides encoding each sgRNA (Table 4) were annealed and inserted into the BbsI site of the pX335 vector (Addgene, 42335), according to protocols available on the Addgene website.

**TABLE 4:**
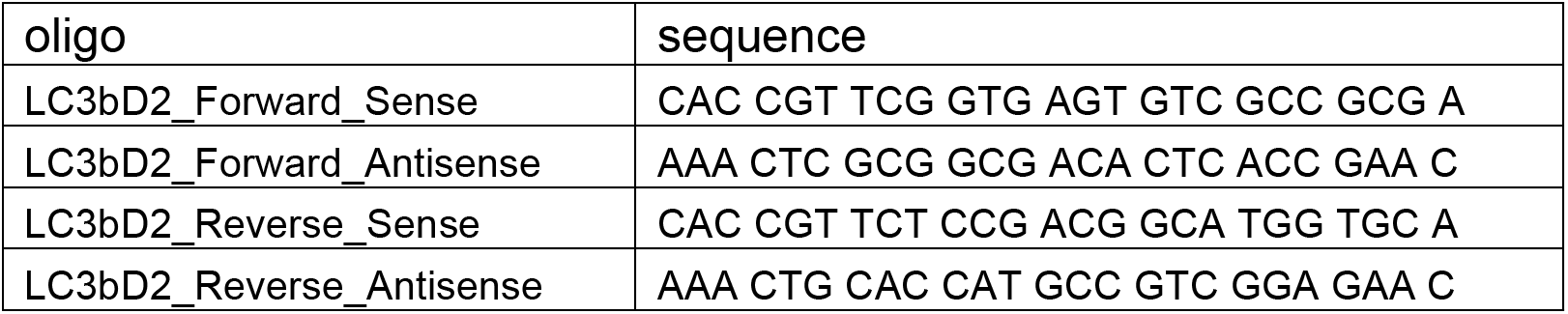
Sense and antisense oligonucleotides cloned into pX335 plasmid

**TABLE 5:** Antibodies used in this study

| **Antibody** | **dilution** | **manufacturer** | **Cat #** |
| --- | --- | --- | --- |
| Rabbit anti-LC3 | 1:1000 | Cell Signaling | 2775S |
| Rabbit anti-ATG5 | 1:1000 | Cell Signaling | 129945 |
| Mouse anti-GAPDH | 1:1000 | Millipore | MAB374 |
| Donkey anti-Mouse iRDye 680RD | 1:10,000 | LICOR | 926-68072 |
| Donkey anti-Rabbit iRDye 800CW | 1:10,000 | LICOR | 926-32213 |

The homology directed repair (HDR) donor vector was synthesized in the pUCminusMCS backbone by Blue Heron Biotechnology (Bothell, WA). The donor sequence consisted of the Dendra2 ORF flanked by 400bp of homologous sequence upstream and downstream of *MAP1LC3B* start codon. 1.25ug of pX335-Forward-sgRNA, 1.25ug of pX335-Reverse-sgRNA, and 2.5ug of the homology donor were transfected into HEK-293T cells using lipofectamine 2000 (Invitrogen, Waltham, MA), according to the manufacturers suggested protocol. Cells were selected based on fluorescence and passaged to homogeneity.

### Western Blotting and antibodies

HEK293T cells were lysed in RIPA buffer (Thermo, Waltham, MA,cat #89900) containing protease inhibitors (Roche, Basel, Switzerland, cat #11836170001).

### siRNA knockdown

HEK293T cells were plated at 60% confluency, then transfected the next day using DharmaFECT 1 Transfection reagent (Dharmacon, LaFayette, CO, T-2001-02) and the following siRNAs from Dharmacon: ON-TARGETplus ATG5 Smartpool siRNA (L-004374-00-0005), ON-TARGETplus MAP1LC3B Smartpool siRNA (L-012846-00-0005) or non-targeting siRNA (D-001810-01-05), at a concentration of either 20nM and 40nM. Cells were imaged and lysates were collected 2d after siRNA transfection.

### Primary screen

HEK293T Dendra2-LC3 cells were plated in HEK complete media at 1.1×10^5^ cells/mL on ViewPlate 384w plates (Perkin Elmer, Waltham, MA, cat #: 6007460) using a Multidrop Combi (Thermo Scientific, cat #: 5840300), adding 50uL to each well. Approximately 48h later, and immediately prior to imaging, HEK complete media was exchanged with Neumo+SOS media using a Biomek FX^P^ laboratory automation workstation (Beckman Coulter, Brea, CA). To avoid dislodging cells during the media exchange, 35uL of the HEK media was removed and replaced with 45uL Neumo+SOS. Another 45uL was removed and replaced with Neumo+SOS, effectively diluting out the concentration of HEK complete media to 6.25%. Cells were imaged with an ImageXpress Micro (Molecular Devices, San Jose, CA) equipped with a 20x objective lens. After imaging in the GFP channel (Semrock, Rochester, NY, FITC-3540B-NTE-ZERO filter) to measure baseline fluorescence intensity, compounds were added using a BioMek FX pintool (Beckman Coulter) at a concentration of 10µM, with the exception of the positive control Torin1, which was added at 1µM. Plates were imaged again 15h after drug addition. One image was acquired per well. Images were background subtracted using FIJI (https://fiji.sc/) with a rolling ball radius of 150. Mean GFP fluorescence intensity was calculated on a whole well basis. Enhancers are defined as: [Sample_15H GFP/0H GFP_] < [DMSO_15H GFP/0H GFP_ - 3SD_DMSO_] and inhibitors as: [Sample_15H GFP/0H GFP_] > [DMSO_15H GFP/0HGFP_ + 3SD_DMSO_]. Using a custom FIJI script that measured the area of occupied by cells, toxic compounds were identified as those that elicited a reduction in cellular area ≥ 3SD, verified by eye.

### Secondary screen

Plating of HEK293T Dendra2-LC3 cells and media exchange were performed as in the primary screen, described above. Two sites per well were imaged in the Texas Red (RFP) (Semrock TxRed-4040C-NTE-ZERO filter) channel prior to photoconversion to establish background fluorescence levels. Photoconversion was accomplished using a 4s DAPI (Semrock Brightline DAPI-5060-NTE-ZERO filter) exposure, and afterwards the cells were immediately imaged once more in the RFP channel. The plate was then removed from the ImageXpress stage and compounds were added to 3 replicate wells via the BioMekFX automation station at a working concentration of 10µM. The plate was then returned to the ImageXpress and imaged every 1.5h in a recurring loop for 16h, while maintaining 5% CO_2_, humidity and a temperature of 37C. Initial optimization studies demonstrated a maximal Z’ = 0.79 within 9h of drug addition, and therefore this time was selected for measuring autophagic flux. We defined enhancers and inhibitors by the following criteria: enhancer, [Sample_9H RFP/(Postconversion RFP - background RFP)_] < [DMSO_9HRFP/(Postconversion RFP - background RFP)_ - 3SD_DMSO_]; inhibitor, [Sample_9H RFP/(Postconversion RFP – background RFP)_] > [DMSO_9H RFP/(Postconversion RFP - background RFP)_+ 3SD_DMSO_]. Images with autofluorescent artifacts were excluded and the remaining images were averaged on a per compound basis.

### RFP-GFP-LC3 assay

HeLa RFP-GFP-LC3 cells^59^ were plated at 80% confluency in HEK complete media. Prior to imaging, cells were switched to Neumo+SOS media. Cells were imaged at baseline as well as 0, 4, 8, and 12h after drug addition. Images were background subtracted using the rolling ball background subtraction plugin in FIJI, with a radius = 150. LC3 puncta were identified and quantified using CellProfiler 3.0 (https://cellprofiler.org/) utilizing a customized pipeline (https://github.com/BarmadaLab/LC3-puncta). Briefly, a series of image processing operations are performed to segment a cell into nuclear and cytoplasmic compartments. The contrast between puncta and background is further enhanced to emphasize LC3 puncta in both GFP and RFP images. Object-oriented colocalization then records the number of cytoplasmic red puncta that are also green, representing autophagosomes.

### Primary neuron survival assay

Primary neurons were dissected from the cortex of embryonic day 20 rat pups and plated at a density of 1×10^5^ cells/well on a laminin/poly-D-lysine coated 96 well plate^686397^ in Neumo complete media (Cell Guidance Systems M07-500). Transfection and longitudinal microscopy were performed as previously described^62^. Briefly, DIV4 neurons were transfected using lipofectamine 2000 (Invitrogen). Neurons were incubated with Lipofectamine-DNA mixtures for 20 minutes followed by washes with Neurobasal media containing 1 mM kynurenic acid to remove residual lipofectamine. Neurons were then placed in half conditioned media and half fresh Neumo+SOS media. A Eclipse Ti inverted microscope (Nikon, Tokyo, Japan) equipped with PerfectFocus, Semrock GFP and TRITC filters, Lambda XL lamp (Sutter Instruments, Novato, CA) and an Andor Zyla 4.2(+) sCMOS camera (Oxford Instruments, Abingdon, UK) was used to acquire images at 20x. To maintain a temperature of 37C and 5% CO2 levels, the microscope was encased in a custom-built plexiglass environmental chamber. Automated stage movements, filter turret rotation, and image acquisition were controlled via µManager with original code written in BeanShell. In-house software was used to assign a barcode for each neuron, measure its fluorescent intensity, and register time of death as described previously^61, 62, 63, 98, 99^. For optical pulse labeling experiments a 1.5s pulse of 405nm light was used for photoconversion. For studies relating the rate of Dendra2-LC3 turnover to neuronal survival only cells that lived the entire duration of OPL imaging were included in the survival analysis.

---

**Figure S1:**
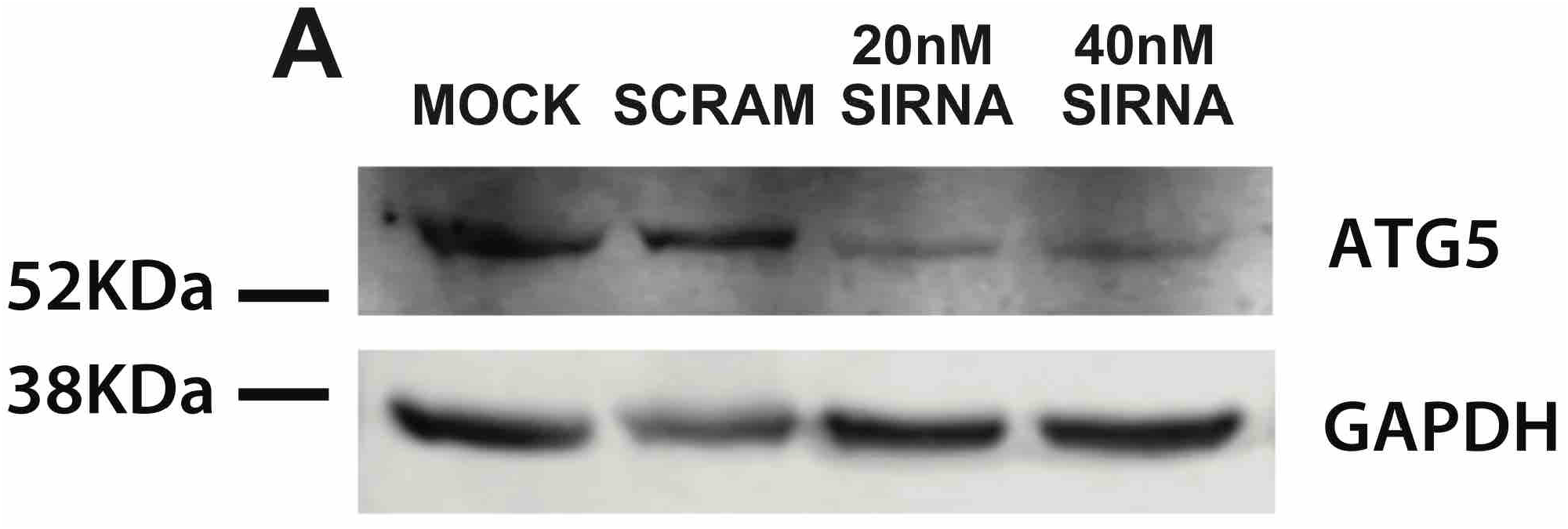
siRNA knockdown of ATG5. HEK293T cells were transfected with 20nM and 40nM ATG5 siRNA, or 40nM non-targeting siRNA. Lysates were collected two days later and blotted with ATG5 and GAPDH antibodies. Relative to scramble siRNA, 20nM and 40nM ATG5 siRNA produced a 58% and 64% knockdown of ATG5 protein, respectively. ATG5 levels were normalized to GAPDH.

**Figure S2:**
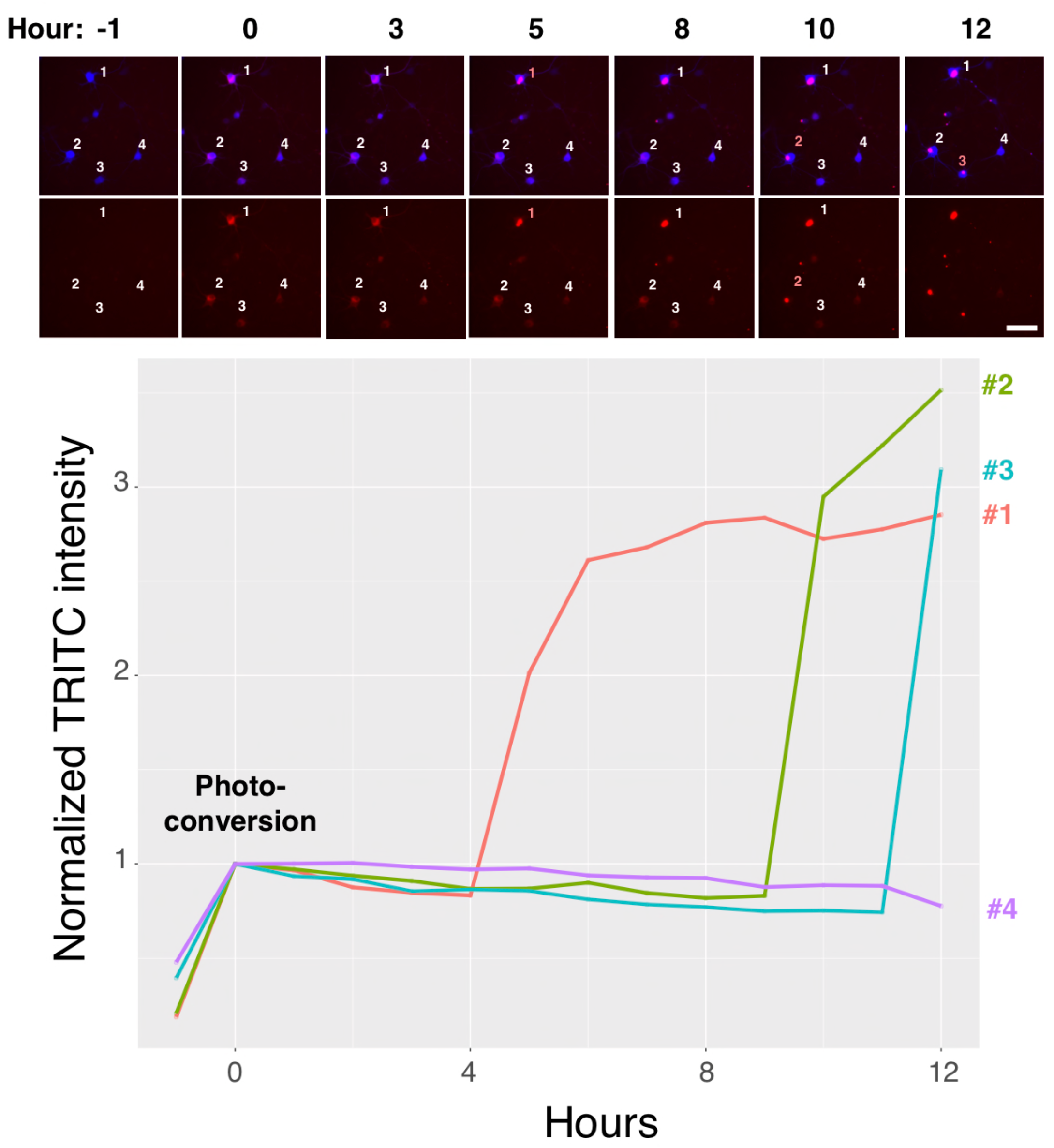
Aggregation of Dendra2 tagged proteins causes a supraphysiological increase in signal. Primary Cortical Neurons were co-transfected on DIV4 with GFP (blue) and a Dendra2-tagged fragment of mutant huntingtin (HTT) carrying a pathologic expansion of 74 glutamine residues (Dendra2-HTT-exon1-Q74)^100^. On the first day following transfection Dendra2-HTT-exon1-Q74 was photoconverted with a 1s pulse of 405nm light and subsequently imaged via automated fluorescence microscopy. The decay in TRITC (red) intensity was measured every hour as a metric for protein degradation. The TRITC intensity spikes upon aggregate formation, preventing accurate measurements of protein half-life. Lines in the plot correspond to the cell labels in white in the image panels. Red cell labels indicate the time point when a cell has formed an aggregate. Scale bar = 50µm

**Figure S3:**
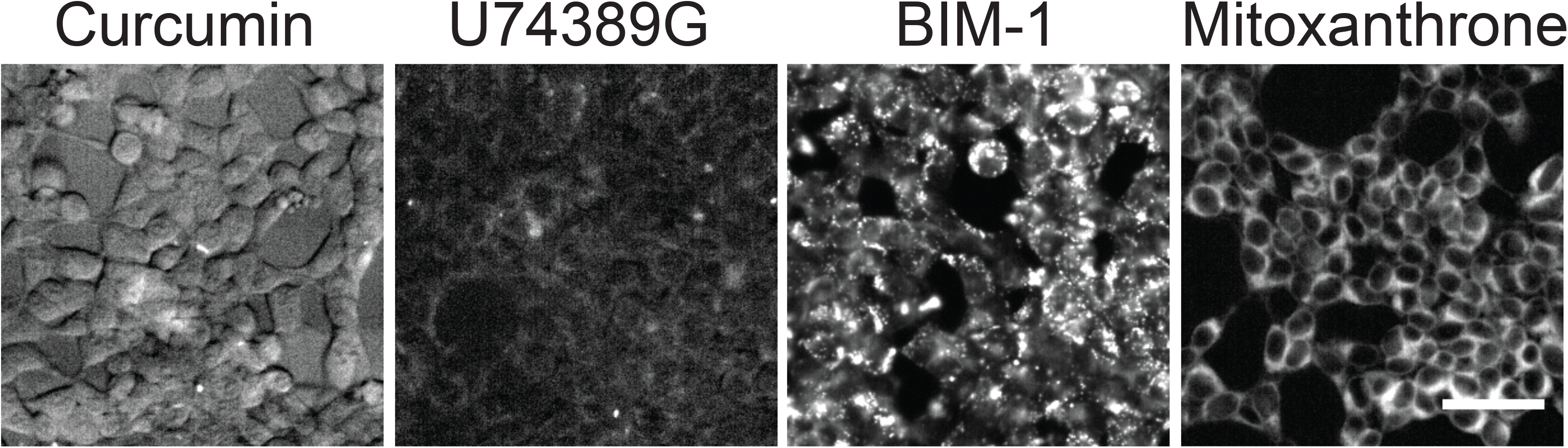
Drugs identified as autophagy modulators display intrinsic fluorescence. Unmodified HEK293T cells imaged in the Texas Red channel 9H after drug treatment. Scale bar = 50µm

**Figure S4:**
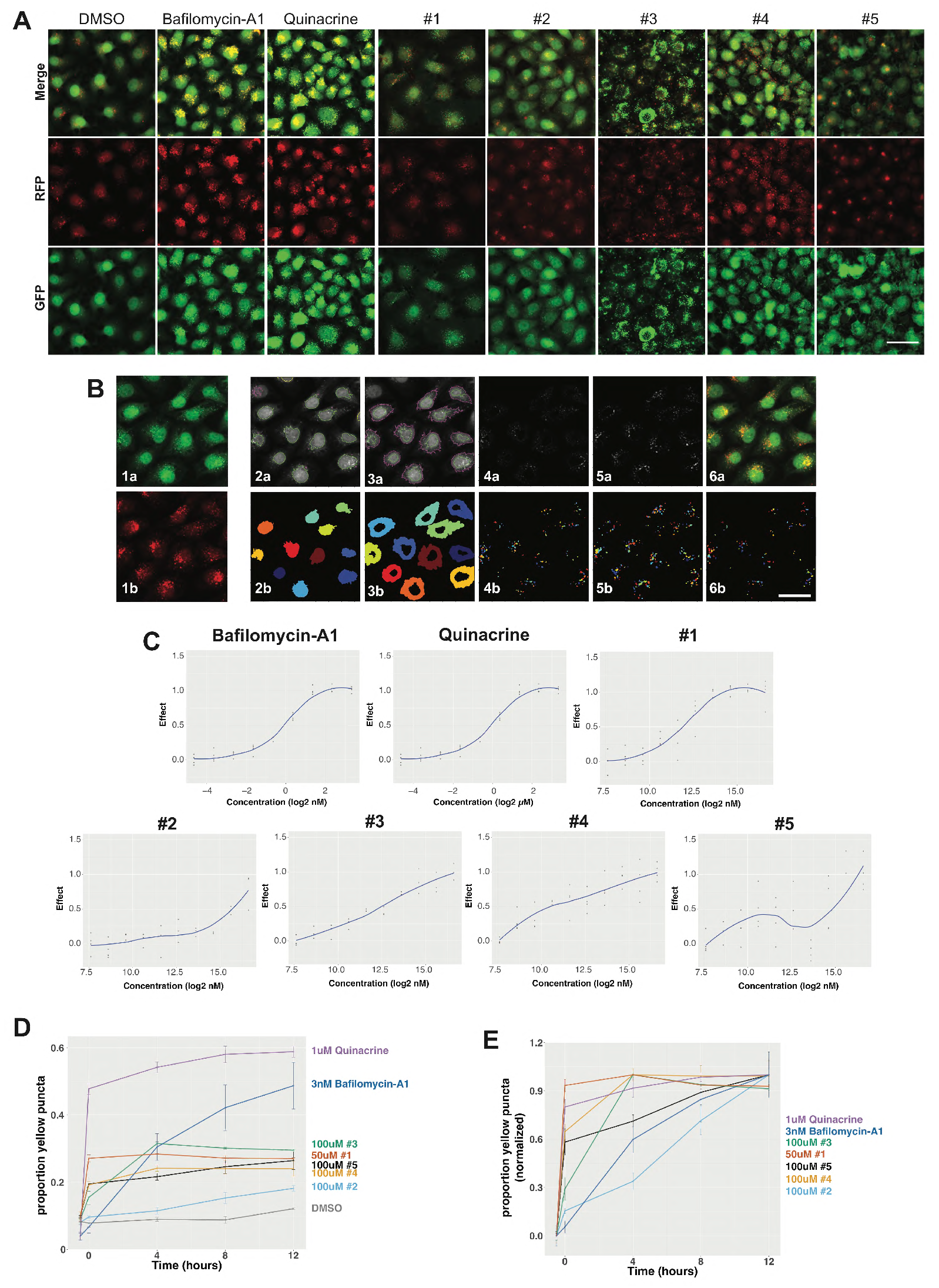
Novel inhibitors block autophagic flux when assessed using independent measures of autophagy. (**A**) HeLa cells expressing a tandem LC3 reporter (RFP-GFP-LC3) were treated with the indicated compounds and imaged 12h later in the RFP (middle panels) and GFP (bottom panels) channels. Composite images are displayed on the top row. Concentrations for each compound (3nM bafilomycin-A1; 1µM Quinacrine; 50µM 245536; and 100µM of 254522, 45808, 234794, and 237373) correspond to the lowest dose resulting in the maximum degree of colocalization between GFP and RFP puncta as calculated in (C). Scale bar: 50µm (**B**) The percentage of RFP(+)/GFP(+) puncta was determined using CellProfiler^101^. Images from the GFP (1a) and RFP (1b) channels are uploaded into CellProfiler, nuclei are identified using the GFP channel (2a-green outlines) and a nuclear mask (2b) is generated. Nuclei that do not pass size or intensity thresholds, or are on the edges of an image, are excluded (2a-purple outlines). Cellular boundaries are identified (3a-purple outlines), and the nuclear mask is subtracted from the newly created cellular mask to produce a cytoplasmic mask (3b). The intensity of cytoplasmic GFP (4a) and RFP (5a) puncta are enhanced, allowing the generation of masks corresponding to puncta in both channels (4b, 5b) and the identification of GFP(+)/RFP(+) autophagosomes (6a, b). Scale bar: 50µm. (**C**) Dose response curves showing the degree of autophagy inhibition with increasing drug concentrations, plotted on a log2 scale. The y-axis represents the proportion of RFP(+) puncta that are GFP(+), with the maximum and minimum set to 1.0 and 0, respectively. Dots represent individual technical replicates. (**D**) RFP-GFP-LC3 HeLa cells were imaged, treated with the lowest dose that produced the maximum response as calculated in (C), then imaged again 0, 4, 8 and 12h after drug treatment. (**E**) Normalized data from (D), depicting the time to maximal effect for each compound. Data in (D, E) represent mean ± SEM from 3 biological replicates, 8 technical replicates each.

**Figure S5:**
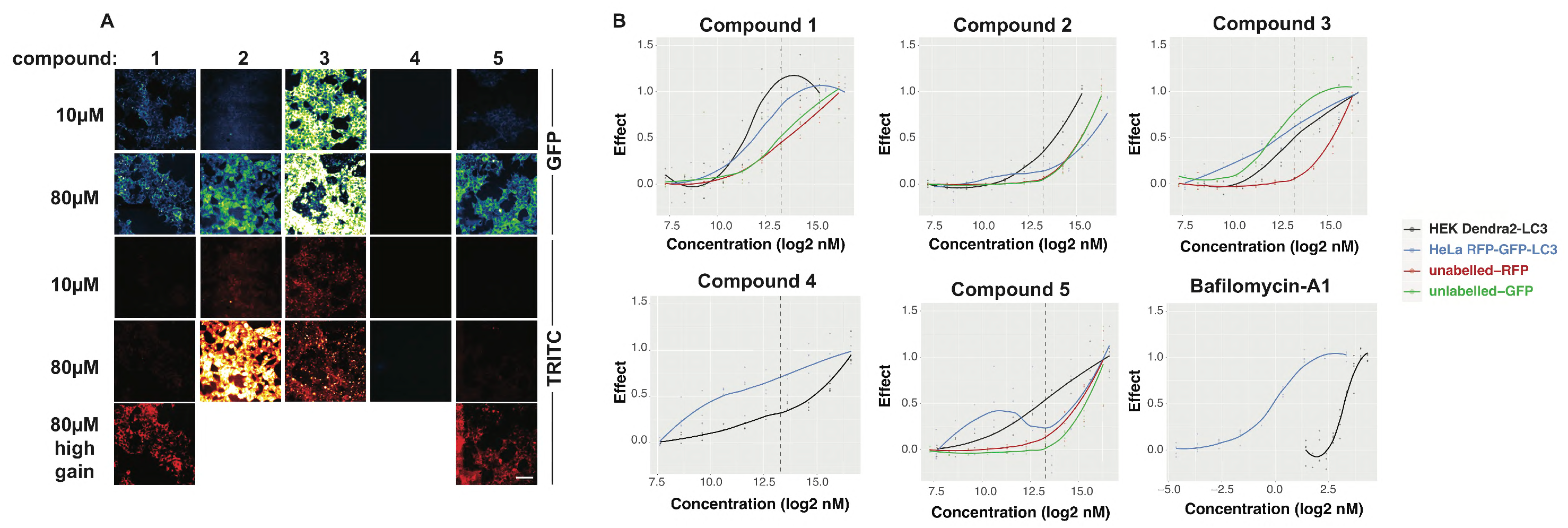
Intrinsic fluorescence confounds flux estimates in a subset of Maybridge library hits. Unmodified HEK293T cells were treated with each compound that registered as an autophagy inhibitor in the screen of the Maybridge 24K library (Fig. 6) Images were acquired in the GFP and TRITC channels 9H after treatment of each drug at either 10µM or 80µM. Images are psuedocolored to accentuate intensity differences. For compounds 1 and 5, 80µM images are presented at higher brightness settings to emphasize low levels of intrinsic fluorescence. All other images are presented at equivalent brightness and contrast. Scale Bar=50µm (**B**) Dose response relationships for each drug in the Dendra-LC3 HEK293T (black), HeLa RFP-GFP-LC3 (blue) autophagic flux assays, as well as intrinsic fluorescence in the GFP (green) and TRITC (red) channels in unmodified HEK cells. In Dendra2-LC3 HEK293T cells the maximum effect represents the greatest increase in RFP intensity 14h after drug treatment. In RFP-GFP-LC3 HeLa cells the effect represents the proportion RFP(+) puncta that are GFP(+), with the maximum and minimum set to 1.0 and 0, respectively. For unlabeled HEK293T cells the maximum fluorescence intensity observed 9H after drug treatment was set to 1 and the lowest to 0. Concentration is plotted in nM on a log(2) scale, with ≥ 3 replicate wells for each concentration shown as colored dots. Dotted vertical lines correspond to 10µM, which is the concentration the compounds were initially screened at.

**Figure S6:**
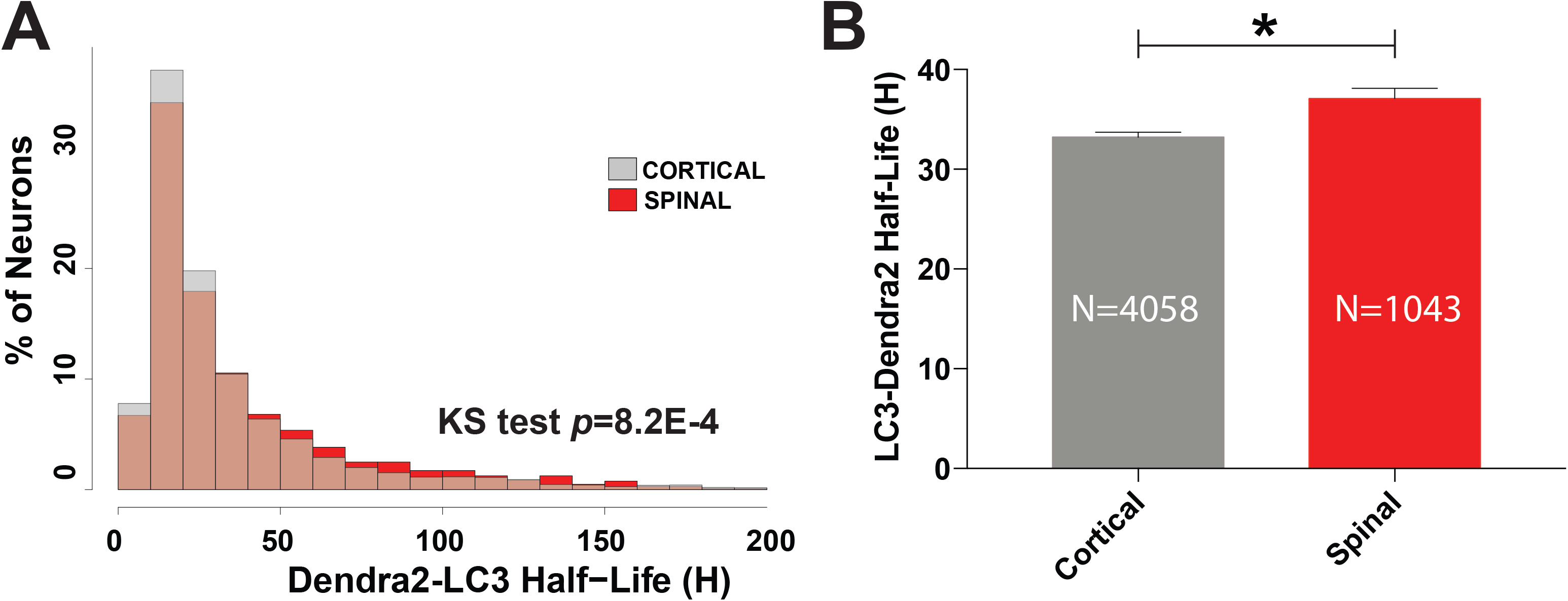
Primary cortical and spinal neurons display subtly different rates of autophagic flux. **A**) Histogram depicting single cell Dendra2-LC3 half-lives in cortical and spinal neurons. Relative to spinal neurons (n=1043), a leftward shift was observed in the Dendra2-LC3 half-life distribution of cortical neurons (n=4058) indicating a greater frequency of cortical neurons exhibiting high rates of autophagic flux compared to spinal neurons (p=8.2×10^-4^, two-sample Kolmogorov-Smirnov (KS) test). (**B**) Consistent with this, the mean single-cell Dendra2-LC3 half-life was slightly reduced in cortical neurons (33.2h) compared to spinal neurons (37.1h) (p=7.1×10^-4^, Welch two sample t-test), indicating higher rates of basal autophagy in cortical neurons.

**Figure 3-source data.**
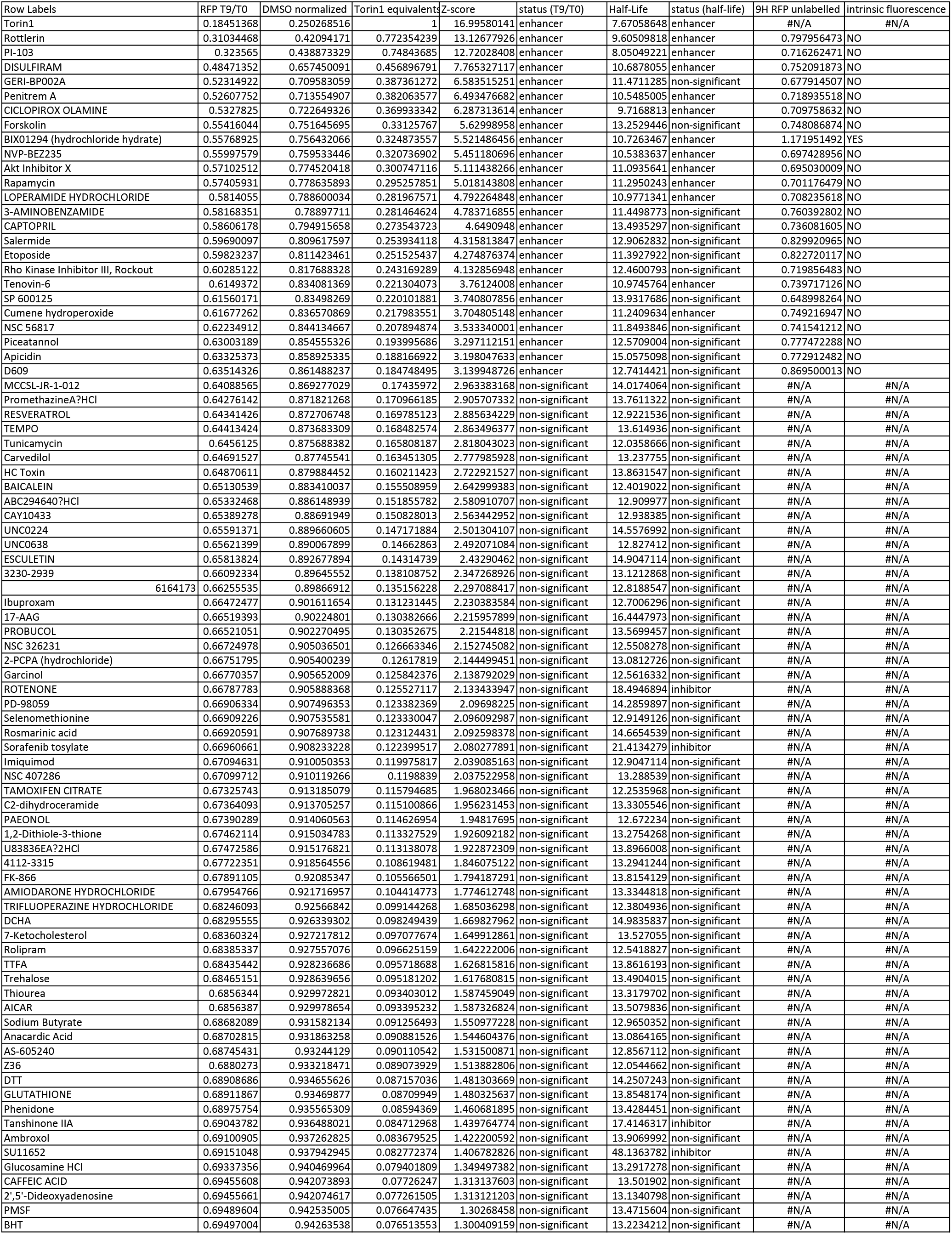
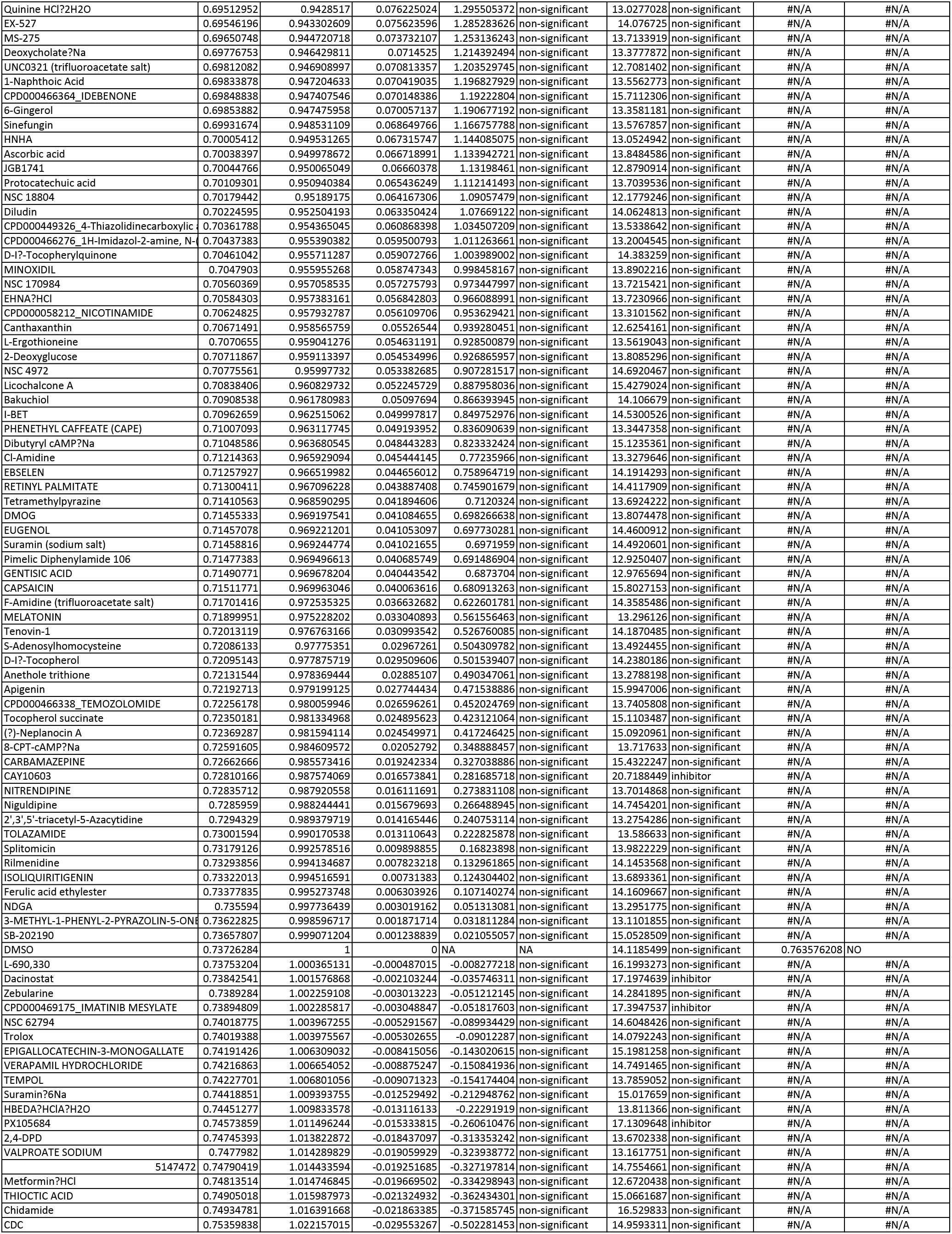
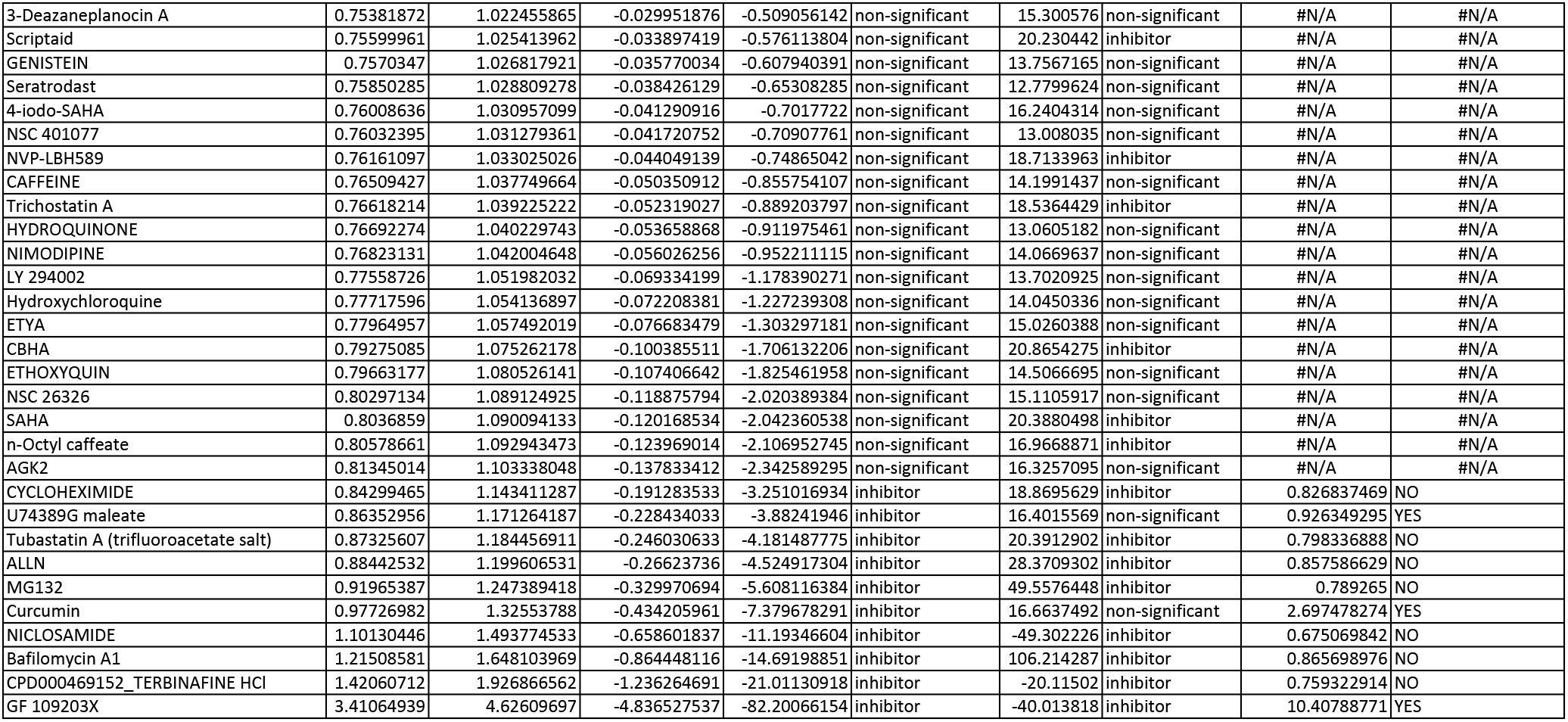

**Figure 5-source data.**
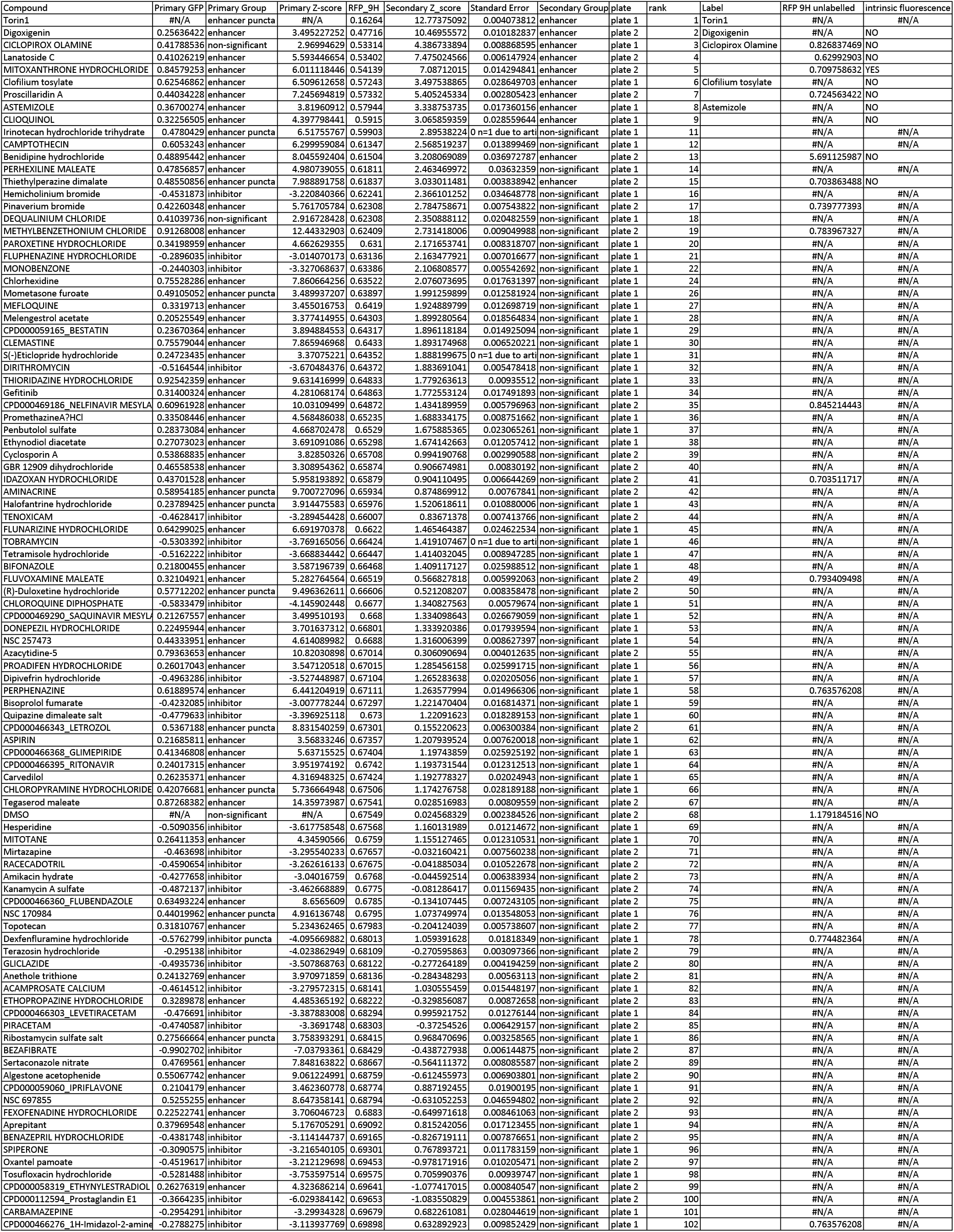
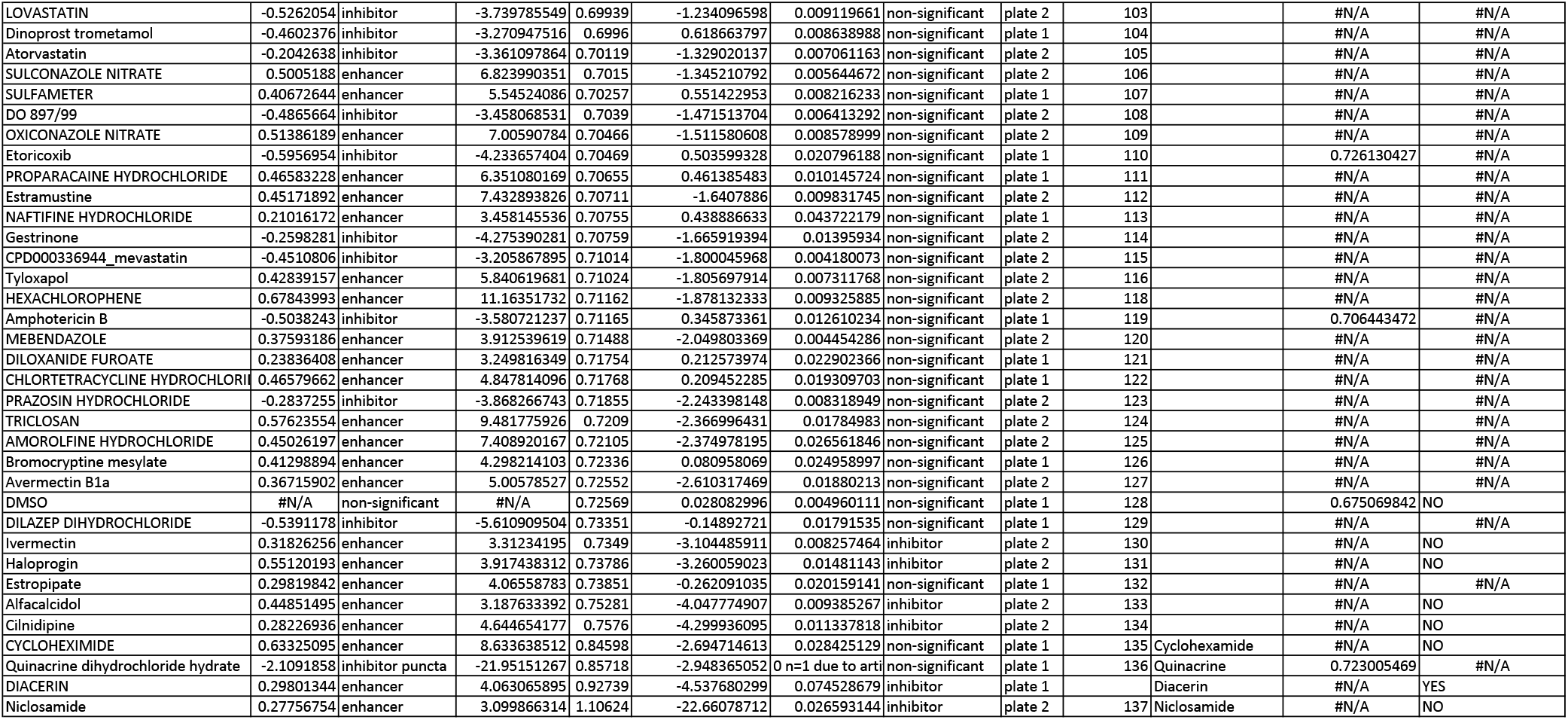

**Supplemental Movie 1: Dendra2-LC3 HEK cells treated with DMSO.**

Dendra2-LC3 HEK cells imaged 6 hours after treatment with DMSO. Presented at 3x actual speed. Scale bar = 10µm

**Supplemental Movie 2: Dendra2-LC3 HEK cells treated with Torin1.**

Dendra2-LC3 HEK cells imaged 6 hours after treatment with 1µM Torin1. Presented at 3x actual speed. Scale bar = 10µm

**Supplemental Movie 3: Dendra2-LC3 HEK cells treated with Bafilomycin-A1.**

Dendra2-LC3 HEK cells imaged 6 hours after treatment with 20nM Bafilomycin-A1. Presented at 3x actual speed. Scale bar = 10µm

## Acknowledgements

This work was supported by the National Institute for Aging (NIA) P30 AG053760, the University of Michigan Protein Folding Disease Initiative, the Center for Chemical Genomics (CCG) at the University of Michigan Life Sciences Institute, Michigan Drug Discovery, and Ann Arbor Active Against ALS. We thank Dr. Vincent Groppi of Michigan Drug Discovery for his continued support and insights; Dr. Lois Weisman and Prof. Tamotsu Yoshimori for RFP-GFP-LC3 HeLa cells; and Dr. Paula Gedraitis (Molecular Devices, LLC) for technical assistance.

## Author Contributions

Conceptualization, N.Safren, S.J.B; Methodology, N.Safren and N.Santoro, Investigation, N.Safren and E.M.T, Writing – Original Draft, N.Safren and S.J.B, Writing – Review and Editing, N.Safren and S.J.B; Funding Acquisition, S.J.B, Resources, S.J.B

## Citations

1. Klionsky DJ. Autophagy as a Regulated Pathway of Cellular Degradation. Science (80-). 2000;290(5497):1717–1721. doi:10.1126/science.290.5497.1717

2. Gatica D, Lahiri V, Klionsky DJ. Cargo recognition and degradation by selective autophagy. 2018. doi:10.1038/s41556-018-0037-z

3. Akematsu T, Akporiaye ET, Al-rubeai M, et al. Guidelines for the use and interpretation of assays for monitoring autophagy (3rd edition). Autophagy. 2016;12(February):1–222. doi:10.1080/15548627.2015.1100356

4. Qu X, Yu J, Bhagat G, et al. Promotion of tumorigenesis by heterozygous disruption of the beclin 1 autophagy gene. J Clin Invest. 2003;112(12):1809–1820. doi:10.1172/JCI20039

5. Yue Z, Jin S, Yang C, Levine AJ, Heintz N. Beclin 1, an autophagy gene essential for early embryonic development, is a haploinsufficient tumor suppressor. Proc Natl Acad Sci. 2003;100(25):15077–15082. doi:10.1073/pnas.2436255100

6. Komatsu M, Waguri S, Ueno T, et al. Impairment of starvation-induced and constitutive autophagy in *Atg7*-deficient mice. J Cell Biol. 2005;169(3):425–434. doi:10.1083/jcb.200412022

7. Galluzzi L, Bravo-San Pedro JM, Levine B, Green DR, Kroemer G. Pharmacological modulation of autophagy: Therapeutic potential and persisting obstacles. Nat Rev Drug Discov. 2017;16(7):487–511. doi:10.1038/nrd.2017.22

8. Rao S, Tortola L, Perlot T, et al. A dual role for autophagy in a murine model of lung cancer. Nat Commun. 2014;5(1):3056. doi:10.1038/ncomms4056

9. Rosenfeldt MT, O’Prey J, Morton JP, et al. p53 status determines the role of autophagy in pancreatic tumour development. Nature. 2013;504(7479):296–300. doi:10.1038/nature12865

10. Galluzzi L, Bravo-San Pedro JM, Demaria S, Formenti SC, Kroemer G. Activating autophagy to potentiate immunogenic chemotherapy and radiation therapy. Nat Rev Clin Oncol. 2017;14(4):247–258. doi:10.1038/nrclinonc.2016.183

11. Guo JY, Xia B, White E. Autophagy-mediated tumor promotion. Cell. 2013;155(6):1216–1219. doi:10.1016/j.cell.2013.11.019

12. Komatsu M, Wang QJ, Holstein GR, et al. Essential role for autophagy protein Atg7 in the maintenance of axonal homeostasis and the prevention of axonal degeneration. Proc Natl Acad Sci U S A. 2007;104(36):14489–14494. doi:10.1073/pnas.0701311104

13. Hara T, Nakamura K, Matsui M, et al. Suppression of basal autophagy in neural cells causes neurodegenerative disease in mice. Nature. 2006;441(7095):885–889. doi:10.1038/nature04724

14. Joo JH, Wang B, Frankel E, et al. The Noncanonical Role of ULK/ATG1 in ER-to-Golgi Trafficking Is Essential for Cellular Homeostasis. Mol Cell. 2016;62(4):491–506. doi:10.1016/J.MOLCEL.2016.04.020

15. Liang C-C, Wang C, Peng X, Gan B, Guan J-L. Neural-specific deletion of FIP200 leads to cerebellar degeneration caused by increased neuronal death and axon degeneration. J Biol Chem. 2010;285(5):3499–3509. doi:10.1074/jbc.M109.072389

16. Nishiyama J, Miura E, Mizushima N, Watanabe M, Yuzaki M. Aberrant Membranes and Double-Membrane Structures Accumulate in the Axons of *Atg5* - Null Purkinje Cells before Neuronal Death. Autophagy. 2007;3(6):591–596. doi:10.4161/auto.4964

17. Menzies FM, Fleming A, Caricasole A, et al. Autophagy and Neurodegeneration: Pathogenic Mechanisms and Therapeutic Opportunities. Neuron. 2017;93(5):1015–1034. doi:10.1016/j.neuron.2017.01.022

18. Pickford F, Masliah E, Britschgi M, et al. The autophagy-related protein beclin 1 shows reduced expression in early Alzheimer disease and regulates amyloid β accumulation in mice. J Clin Invest. May 2008. doi:10.1172/JCI33585

19. Wong E, Cuervo AM. Autophagy gone awry in neurodegenerative diseases. Nat Neurosci. 2010;13(7):805–811. doi:10.1038/nn.2575

20. Ramirez A, Heimbach A, Gründemann J, et al. Hereditary parkinsonism with dementia is caused by mutations in ATP13A2, encoding a lysosomal type 5 P-type ATPase. Nat Genet. 2006;38(10):1184–1191. doi:10.1038/ng1884

21. Kitada T, Asakawa S, Hattori N, et al. Mutations in the parkin gene cause autosomal recessive juvenile parkinsonism. Nature. 1998;392(6676):605–608. doi:10.1038/33416

22. Valente EM, Abou-Sleiman PM, Caputo V, et al. Hereditary early-onset Parkinson’s disease caused by mutations in PINK1. Science. 2004;304(5674):1158–1160. doi:10.1126/science.1096284

23. Tanik SA, Schultheiss CE, Volpicelli-Daley LA, Brunden KR, Lee VMY. Lewy body-like α-synuclein aggregates resist degradation and impair macroautophagy. J Biol Chem. 2013;288(21):15194–15210. doi:10.1074/jbc.M113.457408

24. Martinez-Vicente M, Talloczy Z, Wong E, et al. Cargo recognition failure is responsible for inefficient autophagy in Huntington’s disease. Nat Neurosci. 2010;13(5):567–576. doi:10.1038/nn.2528

25. Ashkenazi A, Bento CF, Ricketts T, et al. Polyglutamine tracts regulate beclin 1-dependent autophagy. Nature. 2017;545(7652):108–111. doi:10.1038/nature22078

26. Wong YC, Holzbaur ELF. The Regulation of Autophagosome Dynamics by Huntingtin and HAP1 Is Disrupted by Expression of Mutant Huntingtin, Leading to Defective Cargo Degradation. J Neurosci. 2014;34(4):1293–1305. doi:10.1523/JNEUROSCI.1870-13.2014

27. Nguyen DKH, Thombre R, Wang J. Autophagy as a common pathway in amyotrophic lateral sclerosis. Neurosci Lett. April 2018. doi:10.1016/J.NEULET.2018.04.006

28. Pottier C, Bieniek KF, Finch N, et al. Whole-genome sequencing reveals important role for TBK1 and OPTN mutations in frontotemporal lobar degeneration without motor neuron disease. Acta Neuropathol. 2015;130(1):77–92. doi:10.1007/s00401-015-1436-x

29. Gijselinck I, Van Mossevelde S, van der Zee J, et al. Loss of *TBK1* is a frequent cause of frontotemporal dementia in a Belgian cohort. Neurology. 2015;85(24):2116–2125. doi:10.1212/WNL.0000000000002220

30. Le Ber I, Camuzat A, Guerreiro R, et al. *SQSTM1* Mutations in French Patients With Frontotemporal Dementia or Frontotemporal Dementia With Amyotrophic Lateral Sclerosis. JAMA Neurol. 2013;70(11):1403–1410. doi:10.1001/jamaneurol.2013.3849

31. Synofzik M, Maetzler W, Grehl T, et al. Screening in ALS and FTD patients reveals 3 novel UBQLN2 mutations outside the PXX domain and a pure FTD phenotype. Neurobiol Aging. 2012;33(12):2949.e13-2949.e17. doi:10.1016/j.neurobiolaging.2012.07.002

32. Teyssou E, Takeda T, Lebon V, et al. Mutations in SQSTM1 encoding p62 in amyotrophic lateral sclerosis: genetics and neuropathology. Acta Neuropathol. 2013;125(4):511–522. doi:10.1007/s00401-013-1090-0

33. Maruyama H, Morino H, Ito H, et al. Mutations of optineurin in amyotrophic lateral sclerosis. Nature. 2010;465(7295):223–226. doi:10.1038/nature08971

34. DeJesus-Hernandez M, Mackenzie IR, Boeve BF, et al. Expanded GGGGCC Hexanucleotide Repeat in Noncoding Region of C9ORF72 Causes Chromosome 9p-Linked FTD and ALS. Neuron. 2011;72(2):245–256. doi:10.1016/J.NEURON.2011.09.011

35. Renton AE, Majounie E, Waite A, et al. A Hexanucleotide Repeat Expansion in C9ORF72 Is the Cause of Chromosome 9p21-Linked ALS-FTD. Neuron. 2011;72(2):257–268. doi:10.1016/j.neuron.2011.09.010

36. Cirulli ET, Lasseigne BN, Petrovski S, et al. Exome sequencing in amyotrophic lateral sclerosis identifies risk genes and pathways. Science (80-). 2015;347(6229):1436–1441. doi:10.1126/SCIENCE.AAA3650

37. Deng H-X, Chen W, Hong S-T, et al. Mutations in UBQLN2 cause dominant X-linked juvenile and adult-onset ALS and ALS/dementia. Nature. 2011;477(7363):211–215. doi:10.1038/nature10353

38. Florey O, Gammoh N, Kim SE, Jiang X, Overholtzer M. V-ATPase and osmotic imbalances activate endolysosomal LC3 lipidation. Autophagy. 2015;11(1):88–99. doi:10.4161/15548627.2014.984277

39. Zoncu R, Bar-Peled L, Efeyan A, Wang S, Sancak Y, Sabatini DM. mTORC1 senses lysosomal amino acids through an inside-out mechanism that requires the vacuolar H(+)-ATPase. Science. 2011;334(6056):678–683. doi:10.1126/science.1207056

40. Barmada SJ, Serio A, Arjun A, et al. Autophagy induction enhances TDP43 turnover and survival in neuronal ALS models. Nat Chem Biol. 2014;10(8):677–685. doi:10.1038/nchembio.1563

41. Chudakov DM, Lukyanov S, Lukyanov KA. Tracking intracellular protein movements using photoswitchable fluorescent proteins PS-CFP2 and Dendra2. Nat Protoc. 2007;2(8):2024–2032. doi:10.1038/nprot.2007.291

42. Meijer AJ, Lorin S, Blommaart EF, Codogno P. Regulation of autophagy by amino acids and MTOR-dependent signal transduction. Amino Acids. 2015;47(10):2037–2063. doi:10.1007/s00726-014-1765-4

43. Wolfson RL, Sabatini DM. Cell Metabolism The Dawn of the Age of Amino Acid Sensors for the mTORC1 Pathway. Cell Metab. 2017;26:301–309. doi:10.1016/j.cmet.2017.07.001

44. Kocaturk NM, Gozuacik D. Crosstalk Between Mammalian Autophagy and the Ubiquitin-Proteasome System. Front Cell Dev Biol. 2018;6:128. doi:10.3389/fcell.2018.00128

45. Mali P, Yang L, Esvelt KM, et al. RNA-guided human genome engineering via Cas9. Science. 2013;339(6121):823–826. doi:10.1126/science.1232033

46. Ran FA, Hsu PD, Lin C-Y, et al. Double nicking by RNA-guided CRISPR Cas9 for enhanced genome editing specificity. Cell. 2013;154(6):1380–1389. doi:10.1016/j.cell.2013.08.021

47. Thoreen CC, Kang SA, Chang JW, et al. An ATP-competitive mammalian target of rapamycin inhibitor reveals rapamycin-resistant functions of mTORC1. J Biol Chem. 2009;284(12):8023–8032. doi:10.1074/jbc.M900301200

48. Maday S, Wallace KE, Holzbaur ELF. Autophagosomes initiate distally and mature during transport toward the cell soma in primary neurons. J Cell Biol. 2012;196(4):407–417. doi:10.1083/jcb.201106120

49. Mizushima N, Yamamoto A, Hatano M, et al. Dissection of autophagosome formation using Apg5-deficient mouse embryonic stem cells. J Cell Biol. 2001;152(4):657–668. http://www.ncbi.nlm.nih.gov/pubmed/11266458. Accessed March 21, 2019.

50. Liu Q, Kang SA, Thoreen CC, et al. Development of ATP-competitive mTOR inhibitors. Methods Mol Biol. 2012;821:447–460. doi:10.1007/978-1-61779-430-8_29

51. Tsvetkov AS, Miller J, Arrasate M, Wong JS, Pleiss MA, Finkbeiner S. A small-molecule scaffold induces autophagy in primary neurons and protects against toxicity in a Huntington disease model. Proc Natl Acad Sci U S A. 2010;107(39):16982–16987. doi:10.1073/pnas.1004498107

52. Balgi AD, Fonseca BD, Donohue E, et al. Screen for chemical modulators of autophagy reveals novel therapeutic inhibitors of mTORC1 signaling. PLoS One. 2009;4(9). doi:10.1371/journal.pone.0007124

53. Akar U, Ozpolat B, Mehta K, Fok J, Kondo Y, Lopez-Berestein G. Tissue transglutaminase inhibits autophagy in pancreatic cancer cells. Mol Cancer Res. 2007;5(3):241–249. doi:10.1158/1541-7786.MCR-06-0229

54. Watanabe-Asano T, Kuma A, Mizushima N. Cycloheximide inhibits starvation-induced autophagy through mTORC1 activation. Biochem Biophys Res Commun. 2014;445(2):334–339. doi:10.1016/j.bbrc.2014.01.180

55. Shakeri A, Cicero AFG, Panahi Y, Mohajeri M, Sahebkar A. Curcumin: A naturally occurring autophagy modulator. J Cell Physiol. 2019;234(5):5643–5654. doi:10.1002/jcp.27404

56. Chignell CF, Bilski P, Reszka KJ, Motten AG, Sik RH, Dahl TA. Spectral and photochemical properties of curcumin. Photochem Photobiol. 1994;59(3):295–302. http://www.ncbi.nlm.nih.gov/pubmed/8016208. Accessed August 6, 2019.

57. Aldrich LN, Goel G, Lassen KG, et al. Small-molecule enhancers of autophagy modulate cellular disease phenotypes suggested by human genetics. Proc Natl Acad Sci. 2015;112(31):E4281–E4287. doi:10.1073/pnas.1512289112

58. Bhaskar K, Timmins GS, Arko-Mensah J, et al. Pharmaceutical screen identifies novel target processes for activation of autophagy with a broad translational potential. Nat Commun. 2015;6(1):1–15. doi:10.1038/ncomms9620

59. Kimura S, Noda T, Yoshimori T. Dissection of the autophagosome maturation process by a novel reporter protein, tandem fluorescent-tagged LC3. Autophagy. 3(5):452–460. http://www.ncbi.nlm.nih.gov/pubmed/17534139. Accessed March 22, 2019.

60. Gupta A, Roy S, Lazar AJF, et al. Autophagy inhibition and antimalarials promote cell death in gastrointestinal stromal tumor (GIST). Proc Natl Acad Sci U S A. 2010;107(32):14333–14338. doi:10.1073/pnas.1000248107

61. Arrasate M, Finkbeiner S. Automated microscope system for determining factors that predict neuronal fate. Proc Natl Acad Sci U S A. 2005;102(10):3840–3845. doi:10.1073/pnas.0409777102

62. Weskamp K, Safren N, Miguez R, Barmada S. Monitoring Neuronal Survival via Longitudinal Fluorescence Microscopy. J Vis Exp. 2019;(143). doi:10.3791/59036

63. Malik AM, Miguez RA, Li X, Ho YS, Feldman EL, Barmada SJ. Matrin 3-dependent neurotoxicity is modified by nucleic acid binding and nucleocytoplasmic localization. Elife. 2018;7:1–30. doi:10.7554/eLife.35977

64. Christensen E. Snecial Articles Multivariate Survival Analysis Using Cox’s Regression Model. Liver. 1987;7(6):1346–1358.

65. Neumann M, Sampathu DM, Kwong LK, et al. Ubiquitinated TDP-43 in frontotemporal lobar degeneration and amyotrophic lateral sclerosis. Science. 2006;314(5796):130–133. doi:10.1126/science.1134108

66. Lee EB, Lee VM-Y, Trojanowski JQ. Gains or losses: molecular mechanisms of TDP43-mediated neurodegeneration. Nat Rev Neurosci. 2012;13(1):38–50. doi:10.1038/nrn3121

67. Barmada SJ, Skibinski G, Korb E, Rao EJ, Wu JY, Finkbeiner S. Cytoplasmic mislocalization of TDP-43 is toxic to neurons and enhanced by a mutation associated with familial amyotrophic lateral sclerosis. J Neurosci. 2010;30(2):639–649. doi:10.1523/JNEUROSCI.4988-09.2010

68. Flores BN, Li X, Malik AM, Martinez J, Beg AA, Barmada SJ. An Intramolecular Salt Bridge Linking TDP43 RNA Binding, Protein Stability, and TDP43-Dependent Neurodegeneration. Cell Rep. 2019;27(4):1133–1150.e8. doi:10.1016/j.celrep.2019.03.093

69. Kodama A, Kaizuka T, Ishihara T, et al. An Autophagic Flux Probe that Releases an Internal Control. Mol Cell. 2016;64(4):835–849. doi:10.1016/j.molcel.2016.09.037

70. Eguchi Y, Makanae K, Hasunuma T, Ishibashi Y, Kito K, Moriya H. Estimating the protein burden limit of yeast cells by measuring the expression limits of glycolytic proteins. Elife. 2018;7:1–3. doi:10.7554/eLife.34595

71. Moriya H. Quantitative nature of overexpression experiments. Mol Biol Cell. 2015;26(22):3932–3939. doi:10.1091/mbc.E15-07-0512

72. Bolognesi B, Lorenzo Gotor N, Dhar R, et al. A Concentration-Dependent Liquid Phase Separation Can Cause Toxicity upon Increased Protein Expression. Cell Rep. 2016;16(1):222–231. doi:10.1016/j.celrep.2016.05.076

73. Rudnick ND, Griffey CJ, Guarnieri P, et al. Distinct roles for motor neuron autophagy early and late in the SOD1 ^G93A^ mouse model of ALS. Proc Natl Acad Sci. 2017;114(39):E8294–E8303. doi:10.1073/pnas.1704294114

74. Panicker LM, Sarkar C, Sgambato JA, et al. Altered TFEB-mediated lysosomal biogenesis in Gaucher disease iPSC-derived neuronal cells. Hum Mol Genet. 2015;24(20):5775–5788. doi:10.1093/hmg/ddv297

75. Leestemaker Y, de Jong A, Witting KF, et al. Proteasome Activation by Small Molecules. Cell Chem Biol. 2017;24(6):725–736.e7. doi:10.1016/J.CHEMBIOL.2017.05.010

76. Tanaka M, Machida Y, Niu S, et al. Trehalose alleviates polyglutamine-mediated pathology in a mouse model of Huntington disease. Nat Med. 2004;10(2):148–154. doi:10.1038/nm985

77. Sarkar S, Davies JE, Huang Z, Tunnacliffe A, Rubinsztein DC. Trehalose, a novel mTOR-independent autophagy enhancer, accelerates the clearance of mutant huntingtin and alpha-synuclein. J Biol Chem. 2007;282(8):5641–5652. doi:10.1074/jbc.M609532200

78. Alers S, Löffler AS, Wesselborg S, Stork B. Role of AMPK-mTOR-Ulk1/2 in the regulation of autophagy: cross talk, shortcuts, and feedbacks. Mol Cell Biol. 2012;32(1):2–11. doi:10.1128/MCB.06159-11

79. Shi W-Y, Xiao D, Wang L, et al. Therapeutic metformin/AMPK activation blocked lymphoma cell growth via inhibition of mTOR pathway and induction of autophagy. Cell Death Dis. 2012;3(3):e275–e275. doi:10.1038/cddis.2012.13

80. Tomic T, Botton T, Cerezo M, et al. Metformin inhibits melanoma development through autophagy and apoptosis mechanisms. Cell Death Dis. 2011;2(9):e199. doi:10.1038/cddis.2011.86

81. Yang H, Peng Y-F, Ni H-M, et al. Basal Autophagy and Feedback Activation of Akt Are Associated with Resistance to Metformin-Induced Inhibition of Hepatic Tumor Cell Growth. Shen H-M, ed. PLoS One. 2015;10(6):e0130953. doi:10.1371/journal.pone.0130953

82. Sarkar S, Floto RA, Berger Z, et al. Lithium induces autophagy by inhibiting inositol monophosphatase. J Cell Biol. 2005;170(7). doi:10.1083/jcb.200504035

83. Zhang L, Wang L, Wang R, et al. Evaluating the Effectiveness of GTM-1, Rapamycin, and Carbamazepine on Autophagy and Alzheimer Disease. 2017;23:801–808. doi:10.12659/MSM.898679

84. Wang I-F, Guo B-S, Liu Y-C, et al. Autophagy activators rescue and alleviate pathogenesis of a mouse model with proteinopathies of the TAR DNA-binding protein 43. Proc Natl Acad Sci. 2012;109(37):15024–15029. doi:10.1073/pnas.1206362109

85. Castillo K, Nassif M, Valenzuela V, et al. Trehalose delays the progression of amyotrophic lateral sclerosis by enhancing autophagy in motoneurons. Autophagy. 2013;9(9):1308–1320. doi:10.4161/auto.25188

86. Sarkar S, Chigurupati S, Raymick J, et al. Neuroprotective effect of the chemical chaperone, trehalose in a chronic MPTP-induced Parkinson’s disease mouse model. Neurotoxicology. 2014;44:250–262. doi:10.1016/j.neuro.2014.07.006

87. Yoon YS, Cho ED, Jung Ahn W, Won Lee K, Lee SJ, Lee HJ. Is trehalose an autophagic inducer? Unraveling the roles of non-reducing disaccharides on autophagic flux and alpha-synuclein aggregation. Cell Death Dis. 2017;8(10):e3091. doi:10.1038/cddis.2017.501

88. Lee HJ, Yoon YS, Lee SJ. Mechanism of neuroprotection by trehalose: Controversy surrounding autophagy induction. Cell Death Dis. 2018;9(7). doi:10.1038/s41419-018-0749-9

89. Yarasheski KE, Davidson NO, Heitmeier MR, et al. Trehalose inhibits solute carrier 2A (SLC2A) proteins to induce autophagy and prevent hepatic steatosis. Sci Signal. 2016;9(416):ra21–ra21. doi:10.1126/scisignal.aac5472

90. Bellozi PMQ, Lima IV de A, Dória JG, et al. Neuroprotective effects of the anticancer drug NVP-BEZ235 (dactolisib) on amyloid-β 1-42 induced neurotoxicity and memory impairment. Sci Rep. 2016;6(May):25226. doi:10.1038/srep25226

91. Evans CS, Holzbaur ELF. Autophagy and mitophagy in ALS. Neurobiol Dis. 2019;122:35–40. doi:10.1016/j.nbd.2018.07.005

92. Valenzuela V, Nassif M, Hetz C. Unraveling the role of motoneuron autophagy in ALS. Autophagy. 2018;14(4):733–737. doi:10.1080/15548627.2018.1432327

93. Zhang X, Li L, Chen S, et al. Rapamycin treatment augments motor neuron degeneration in SOD1 G93A mouse model of amyotrophic lateral sclerosis) Rapamycin treatment augments motor neuron degeneration in SOD1 G93A mouse model of amyotrophic lateral sclerosis. Autophagy. 2011;7(4):412–425. doi:10.4161/auto.7.4.14541

94. Fox JH, Connor T, Chopra V, et al. The mTOR kinase inhibitor Everolimus decreases S6 kinase phosphorylation but fails to reduce mutant huntingtin levels in brain and is not neuroprotective in the R6/2 mouse model of Huntington’s disease. Mol Neurodegener. 2010;5(1):26. doi:10.1186/1750-1326-5-26

95. Maday S, Holzbaur ELF. Compartment-Specific Regulation of Autophagy in Primary Neurons. J Neurosci. 2016;36(22):5933–5945. doi:10.1523/jneurosci.4401-15.2016

96. Subach OM, Patterson GH, Ting L-M, Wang Y, Condeelis JS, Verkhusha V V. A photoswitchable orange-to-far-red fluorescent protein, PSmOrange. Nat Methods. 2011;8(9):771–777. doi:10.1038/nmeth.1664

97. Archbold HC, Jackson KL, Arora A, et al. TDP43 nuclear export and neurodegeneration in models of amyotrophic lateral sclerosis and frontotemporal dementia. Sci Rep. 2018;8(1):4606. doi:10.1038/s41598-018-22858-w

98. Barmada SJ, Ju S, Arjun A, et al. Amelioration of toxicity in neuronal models of amyotrophic lateral sclerosis by hUPF1. Proc Natl Acad Sci. 2015;112(25):7821–7826. doi:10.1073/pnas.1509744112

99. Sharkey LM, Safren N, Pithadia AS, et al. Mutant UBQLN2 promotes toxicity by modulating intrinsic self-assembly. Proc Natl Acad Sci. 2018;115(44):E10495–E10504. doi:10.1073/pnas.1810522115

100. Tsvetkov AS, Arrasate M, Barmada S, et al. Proteostasis of polyglutamine varies among neurons and predicts neurodegeneration. Nat Chem Biol. 2013;9(9):586–592. doi:10.1038/nchembio.1308

101. McQuin C, Goodman A, Chernyshev V, et al. CellProfiler 3.0: Next-generation image processing for biology. Misteli T, ed. PLoS Biol. 2018;16(7):e2005970. doi:10.1371/journal.pbio.2005970

